# Myeloid cell reprogramming drives enhanced defense against *Streptococcus pneumoniae* lung infection following exposure to commensal *Prevotella*

**DOI:** 10.64898/2026.06.25.734552

**Authors:** Sara N. Stoner, Eric D. Larson, Sam Fulte, Steven C. Shaw, Erin R. Fish, Edward N. Janoff, Matthias Mack, Sarah E. Clark

## Abstract

Clinical data link the prevalent respiratory tract anaerobe *Prevotella* with reduced pneumonia mortality, but the mechanisms directing *Prevotella* regulation of lung immune homeostasis are unclear. Here, single-cell RNA sequencing was employed to define the transcriptional immune signatures underlying improved clearance of *Streptococcus pneumoniae* following lung exposure to *Prevotella melaninogenica*. Overall, we observed a substantial shift in myeloid cell transcriptional programming from interferon-dominant to a more antibacterial profile in *S. pneumoniae*-infected mice after pre-exposure to *P. melaninogenica*, correlating with increased macrophage and neutrophil phagocytosis of *S. pneumoniae* and improved pathogen clearance. In neutrophils, TNF signaling through TNFR2 was essential for increased antimicrobial function. Moreover, improved defense required CCR2-dependent monocyte-derived macrophages, with selective enrichment of more a mature Cxcl3+ population which was distinct from the hallmark *S. pneumoniae*-associated C1qa+ population enriched in the absence of effective clearance. Together, these findings inform the myeloid cell transcriptional changes associated with natural infection resistance mediated by pulmonary microbial exposures.

## Introduction

*Streptococcus pneumoniae* (the pneumococcus) is an opportunistic pathogen and a leading cause of community-acquired pneumonia worldwide ^1,2^. While carriage rates vary by location, up to 65% of children and 10% of adults are asymptomatically colonized in the nasopharynx ^3–5^. In a subset of colonized individuals, *S. pneumoniae* is aspirated into the lungs, resulting in pneumococcal pneumonia. The factors that determine host resilience versus progression to infection remain poorly understood.

Growing evidence points to the respiratory tract microbiome as a key determinant of pneumonia outcomes ^6–8^. The upper respiratory tract acts as a microbial reservoir, with commensal bacteria continuously aspirated into the lung, particularly those from the oropharynx, which serves as the primary source of transient microbial exposures in the lower airway ^9,10^. Greater microbial diversity in the upper and lower airway correlates with improved survival in pneumonia patients ^11–13^, whereas depletion of obligate anaerobes, including oral commensals ^11^, through antibiotic treatment or chronic oxygen therapy is linked to worse outcomes ^14^.

*Prevotella* species are core members of the oral microbiome and are consistently among the most abundant bacterial taxa detected in the lungs of healthy individuals ^9,15^. Higher *Prevotella* abundance is inversely associated with respiratory pathogens such as *S. pneumoniae* in both the upper airway ^13,16^ and the lungs ^9,17,18^. Specifically, the species *Prevotella melaninogenica* is enriched in healthy individuals compared to patients suffering from pneumonia, acute respiratory distress syndrome (ARDS), and exacerbations of chronic obstructive pulmonary disease (COPD) ^19–21^. In one report, *P. melaninogenica* was the most discriminatory species distinguishing healthy subjects from those with pneumococcal pneumonia ^13^. Beyond correlations with health, *Prevotella* are increasingly recognized as immunomodulatory. Bronchoalveolar lavage samples enriched with *Prevotella* exhibit increased numbers of activated neutrophils and higher Th17 cell frequencies ^22,23^, and lung transplant recipients with more diverse microbial profiles containing *Prevotella* have elevated expression of IL-10 ^24^. Additionally, a single dose of *P. melaninogenica*, in combination with two other oral commensals, increased airway clearance of *S. pneumoniae* from the lungs of mice at 24 hours and up to 14 days post-exposure to the commensal mixture^25^. These findings suggest that *Prevotella* may shape lung immune tone and infection susceptibility.

To mechanistically address the link between *Prevotella* and improved pneumonia outcomes, we previously determined that administration of live or heat-killed *P. melaninogenica* 24 hours prior to a *S. pneumoniae* infection promoted rapid clearance of *S. pneumoniae* from the lungs, which required neutrophils^26^. While several airway *Prevotella* species were similarly protective, we also identified a non-protective *Prevotella* species, indicating a unique, species-specific mechanism of enhanced immune defense^26^. Further, we demonstrated that *P. melaninogenica* exposure improved lung clearance of *Staphylococcus aureus*, demonstrating protective function against multiple respiratory pathogens^27^. Single-cell RNA sequencing (scRNA-seq) has emerged as a powerful tool to resolve the heterogeneity of tissue immune populations in health and disease, though the signaling programs governing immune defense during bacterial pneumonia remain poorly defined. Here, we used scRNA-seq to define the transcriptional immune landscape during *S. pneumoniae* in susceptible versus resistant hosts, with resistance modeled by improved pathogen clearance following *P. melaninogenica* priming. Transcriptional analysis revealed significant changes in myeloid cells, which were dominated by interferon signaling during *S. pneumoniae* infection in susceptible mice but shifted to more pro-phagocytic and antibacterial defense responses in *Prevotella*-exposed mice with enhanced pathogen clearance. The consequences of altered myeloid cell reprogramming were defined by evaluating innate immune cell recruitment to the lungs and the functional activity of neutrophil and macrophage populations. We identified a critical role for neutrophil-intrinsic TNFα signaling using *Tnf*^-/-^, *Tnfrsf1a*^-/-^ and *Tnfrsf1b*^-/-^ mice, revealing a selective requirement for TNFR2 signaling to license neutrophil serine protease activity. Finally, we uncovered a unique population of Cxcl3+ monocyte-derived macrophages associated with altered transcriptional programming that differed from the less mature C1qa+ population enriched during *S. pneumoniae* infection in susceptible mice. Overall, recruited monocytes were required for *Prevotella*-enhanced defense, confirmed by improved macrophage bacterial phagocytosis and loss of protection in CCR2-depleted mice. Together, this study defines a commensal-driven innate immune program in the lung that is mechanistically distinct from canonical pneumococcal inflammation and improves baseline protection during acute bacterial pneumonia.

## Results

### *P. melaninogenica* exposure alters the myeloid cell response to pneumococcal pneumonia

To model the effects of *P. melaninogenica* on the immune response to acute *S. pneumoniae* lung infection, mice were intratracheally instilled with heat-killed (HK) *P. melaninogenica* 24-hours prior to *S. pneumoniae* infection. Mice primed with HK *P. melaninogenica* had significantly reduced *S. pneumoniae* burdens at 24 hours post-infection compared to unprimed mice, with no detectable *S. pneumoniae* infection in two-thirds of the *P. melaninogenica*-exposed mice (**Fig. 1A**). Single-cell RNA sequencing was performed on lung tissue of naïve mice, *P. melaninogenica*-exposed mice, and *S. pneumoniae*-infected mice with or without prior exposure to *P. melaninogenica*. Naïve and *P. melaninogenica*-exposed mice were compared to further define the immune response to *P. melaninogenica* in the absence of *S. pneumoniae* infection. Additionally, *S. pneumoniae*-infected mice pre-exposed to *P. melaninogenica* or phosphate-buffered saline (PBS) were compared to characterize transcriptional changes in the setting of susceptibility versus enhanced defense against pneumococcal challenge (**Fig. 1B**). Clustering analysis identified 30 distinct cell populations in lung tissue, including myeloid, adaptive, epithelial and endothelial cells (**Fig. S1A-D**). Myeloid cell populations included monocytes, macrophages, neutrophils, dendritic cells, and basophils (**Fig. 1C-D**). Broad changes among several myeloid cell populations were apparent in response to both *S. pneumoniae* infection and *P. melaninogenica* exposure (**Fig. 1E**). Two neutrophil populations were detected, both of which expanded in *P. melaninogenica*-exposed and *S. pneumoniae*-infected mice compared to naïve mice. Recruited immune cells detected at baseline in naïve mice are likely in the marginated vascular compartment of the lungs, as reported elsewhere^28^. Neutrophil recruitment increased in both *P. melaninogenica*-exposed and *S. pneumoniae*-infected mice, but was lower in mice pre-exposed to *P. melaninogenica* with enhanced *S. pneumoniae* clearance (**Fig. 1F**). Inflammatory monocytes were recruited to the lung tissue in *P. melaninogenica*-exposed mice and in even greater numbers in *S. pneumoniae*-infected mice (**Fig. 1F**). As with neutrophils, numbers of recruited inflammatory monocytes were lower with dual-bacterial exposure than with either alone. Tissue-resident alveolar macrophages (AMs), prominent at baseline, were depleted in both *P. melaninogenica*-exposed and *S. pneumoniae*-infected mice, alone and in combination, as anticipated following an acute inflammatory stimulus ^29,30^, but more subtly with *P. melaninogenica* alone (**Fig. 1F**). A minority population of macrophages with high expression of cell division genes was present in all conditions (**Fig. S2**), consistent with prior reports of a proliferative AM subpopulation ^30^.

**Figure 1:**
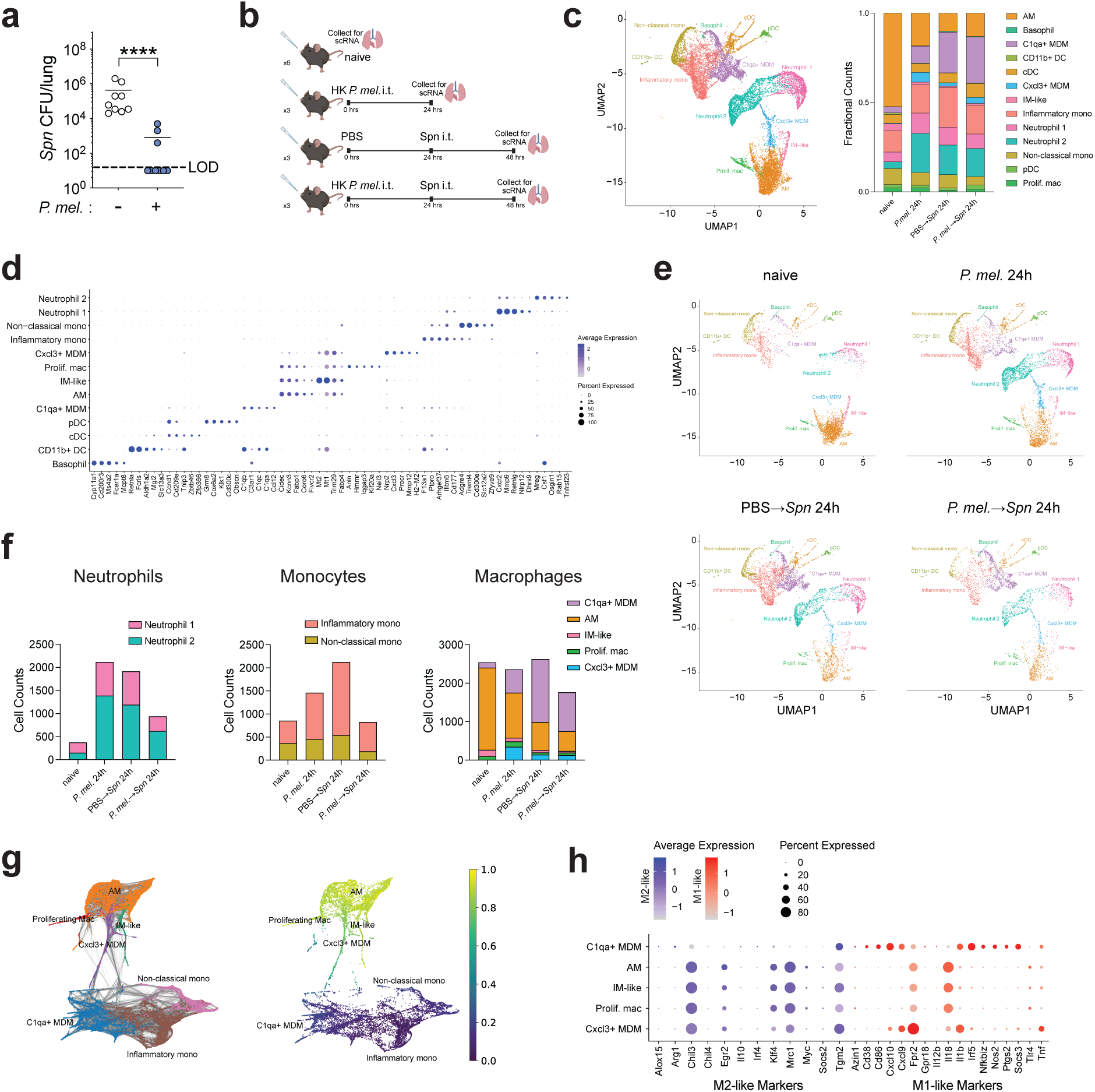
*P. melaninogenica* exposure alters the myeloid cell response to pneumococcal pneumonia. **(A)** *Spn* burdens in lung tissue of mice with exposure to PBS or *P. mel* HK 24 hours prior to infection (mean ± SEM; n = 9 biological replicates pooled from 3 independent experiments). (**B**) Workflow of scRNA-seq experiment sample collection, with n = 6 mice for naïve group and n = 3 mice for all other groups. (**C**) UMAP projection of myeloid cell clusters from all 4 conditions (naïve, *P. mel* 24h, PBS->*Spn* 24h, *P. mel*->*Spn* 24h) and bar plot displaying relative proportions of myeloid populations in each condition. (**D**) Dot plot showing top 5 differentially expressed genes (DEGs) in myeloid cell clusters. (**E**) UMAP projections of myeloid cell clusters split by condition. (**F**) Cell counts of neutrophil, monocytes and macrophage populations detected in scRNA-seq samples by condition. (**G**) ForceAtlas2 ^90^ projection of monocyte and macrophage populations (left) and Partition-based graph abstraction (PAGA)-inferred pseudotime values (right). (**H**) Expression of M1-like and M2-like marker genes across macrophage populations. Abbreviations: AM = alveolar macrophage, IM = interstitial macrophage, MDM = monocyte-derived macrophage, mono = monocyte, mac = macrophage, DC = dendritic cell, cDC = conventional DC, pDC = plasmacytoid DC. Data were analyzed by Mann-Whitney test (**A**), \*\*\*\**p*<0.0001.

Of note, two distinct populations of monocyte-derived macrophages (MDM) were identified which differed proportionally between groups; a prominent C1qa+ MDM population selectively enriched in *S. pneumoniae*-infected mice and a smaller Cxcl3+ MDM population predominantly enriched in *P. melaninogenica*-exposed mice (**Fig. 1F**). Partition-based graph abstraction (PAGA) clustering and pseudotime analysis revealed that C1qa+ MDMs clustered closely with immature monocytes, whereas Cxcl3+ MDMs clustered closer to the mature, tissue resident-like macrophages (**Fig. 1G**). Additionally, the prominent C1qa+ MDMs expressed high levels of pro-inflammatory “M1-like” signature genes, while the Cxcl3+ MDMs and tissue resident-like macrophage populations showed greater expression of pro-resolution “M2-like” signature genes (**Fig. 1H**). This pattern aligns with reports indicating that recruited macrophage populations exhibit an M1-like phenotype, and tissue-resident macrophages are more M2-like during acute lung injury ^30,31^. These findings highlight a key distinction in the type and maturation state of MDMs recruited to the lungs during *S. pneumoniae* infection, associated with impaired bacterial clearance, compared to the MDMs recruited in response to *P. melaninogenica*.

Given our prior finding that toll-like receptor 2 (TLR2) signaling was required for *P. melaninogenica-*induced neutrophil activation and lung clearance of *S. pneumoniae* ^26^, we examined *Tlr2* expression patterns. *Tlr2* was primarily expressed in myeloid cells (**Fig. S3A**), focusing our analysis on myeloid cell types to identify gene expression pathways associated with *P. melaninogenica*-mediated protection. Unbiased pathway enrichment analysis of *S. pneumoniae*-infected mice with versus without *P. melaninogenica* priming revealed distinctly enriched immune activation pathways, including TNF signaling and antigen presentation, in myeloid cells (**Fig. S3B**). Differential signaling pathway enrichment was also apparent among other non-myeloid cell types including epithelial cells and adaptive immune cells (**Fig. S3C**). Together, these findings identify distinct differences in myeloid cell populations and gene expression patterns in the lungs of mice infected with *S. pneumoniae* following pre- exposure to *P. melaninogenica,* when improved *S. pneumoniae* clearance was observed.

### Immune priming by *P. melaninogenica* induces a coordinated recruitment of myeloid cells

Given the large population and gene expression pattern changes associated with *P. melaninogenica* exposure, we next examined myeloid cell recruitment over time in *P. melaninogenica*-primed mice via flow cytometry (**Fig. S4A**). Neutrophils and Ly6C+ monocytes and MDMs were recruited to the lung tissue as early as 6 hours post-exposure to *P. melaninogenica*, with persistence at 24 hours (**Fig. 2A**). This cellular recruitment coincided with elevated concentrations of neutrophil-recruiting chemokines CXCL1, CXCL2 and CXCL5 as well as the monocyte chemoattractant CCL2 in lung bronchoalveolar lavage fluid (**Fig. 2B**). CellChat analysis, which predicts cell-cell communication networks, in all treatment groups combined revealed multiple significantly enriched chemokine signaling pathways. For neutrophil-recruiting chemokines, *Cxcl2-Cxcr2* and *Cxcl3-Cxcr2* signaling pathways were both significantly enriched, with the Cxcl3+ MDM population predicted as a major source of CXCL2 sensed by Cxcr2-expressing neutrophils. CellChat analysis also identified the *Ccl6-Ccr1* signaling pathway as significantly enriched, with several macrophage populations predicted to signal to Ccr1-expressing neutrophils (**Fig. 2C-E**). *Cxcl1* signaling networks were not significantly enriched. Stratification by condition showed that *Cxcl2*, *Cxcl3*, and *Ccl6* were highly expressed in macrophage populations from *P. melaninogenica*-exposed mice, particularly the Cxcl3+ MDMs which were selectively enriched in this group, consistent with the high neutrophil cell counts observed in *P. melaninogenica*-exposed mice. These data highlight the transcriptional and soluble mechanisms underlying macrophage-mediated recruitment of neutrophils and monocytes in response to *P. melaninogenica* exposure.

**Figure 2:**
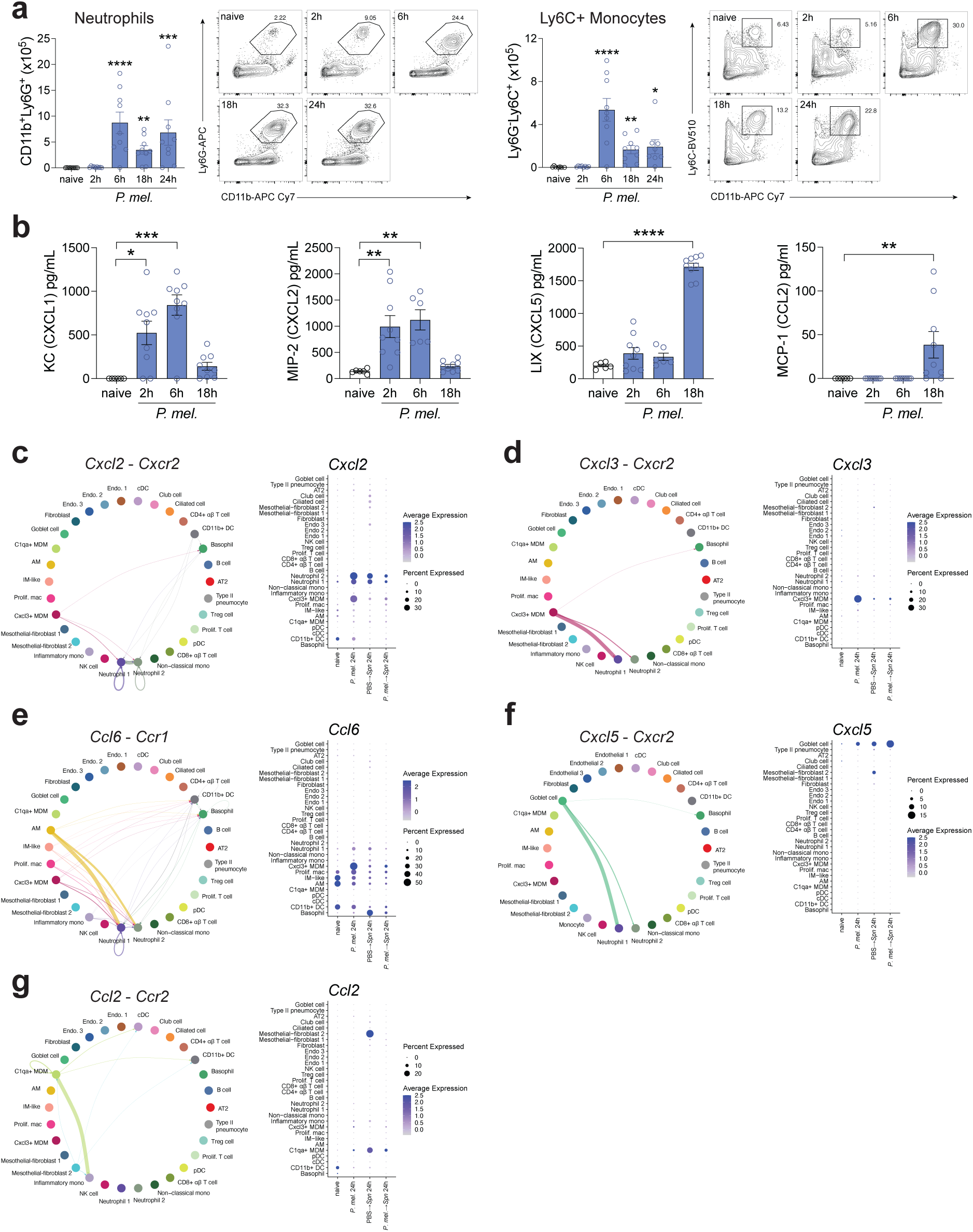
Immune priming by *P. melaninogenica* induces a coordinated recruitment of myeloid cells. (**A**) Quantification of CD11b+Ly6G+ neutrophils and Ly6G-Ly6C+ monocytes/macrophages in the lung tissue via flow cytometry after 0, 2, 6, 18, or 24 hours of i.t. exposure to *P. mel* HK (mean ± SEM; n = 8-9 biological replicates pooled from 3 independent experiments). (**B**) Detection of CXCL1, CXCL2, CXCL5, and CCL2 chemokines in lung bronchoalveolar lavage (BAL) after 0, 2, 6, 18, or 24 hours of i.t. exposure to *P. mel* HK (mean ± SEM; n = 6-9 biological replicates pooled from 3 independent experiments). CellChat circle plots displaying inferred cell-cell communication networks for (**C**) *Cxcl2*-*Cxcr2*, (**D**) *Cxcl3*-*Cxcr2*, (**E**) *Ccl6*-*Ccr1*, (**F**) *Cxcl5*-*Cxcr2*, and (**G**) *Ccr2*-*Ccl2* signaling along with dot plots indicating expression of corresponding chemokine genes by condition. Data were analyzed by Kruskal-Wallis with Dunn’s post hoc test (**A,B**) or ordinary one-way ANOVA with Dunnett’s post hoc test (**B**), \**p*<0.05, \*\**p*<0.01, \*\*\**p*<0.001, \*\*\*\**p*<0.0001.

In addition to macrophages, other cell populations were predicted to contribute to myeloid cell recruitment based on CellChat analysis. Among these, neutrophils were predicted to engage in autocrine signaling, as they expressed both chemokines and their corresponding receptors (**Fig. 2C**), suggesting that neutrophils contribute to their own recruitment through positive feedback. *Cxcl5*, which encodes a neutrophil-recruiting chemokine produced by epithelial cells, was expressed by goblet cells and predicted to signal to *Cxcr2*-expressing neutrophils (**Fig. 2F**). Lastly, *Ccl2*, which encodes a monocyte-recruiting chemokine, was expressed by fibroblasts and C1qa+ MDMs, and was predicted to signal to *Ccr2*-expressing inflammatory monocytes (**Fig. 2G**). Condition-specific analysis revealed that fibroblasts from *S. pneumoniae*-infected mice expressed high levels of *Ccl2* (**Fig. 2G**), correlating with the increased inflammatory monocyte counts observed in these mice compared to other groups. Both the C1qa+ MDMs and Cxcl3+ MDMs also expressed the T cell-recruiting chemokines *Cxcl16*, *Cxcl10*, and *Cxcl9*, predicted to signal to *Cxcr6* and *Cxcr3*-expressing regulatory T cells (**Fig. S4B-D**). Together, these data highlight a coordinated signaling program with predicted contributions from macrophages and epithelial cells to the recruitment of innate immune cells, particularly neutrophils and monocytes, to the lungs following *P. melaninogenica* exposure.

### *P. melaninogenica* induces neutrophil transcriptional profiles associated with enhanced phagocytosis and killing of *S. pneumoniae*

We previously reported the unique capacity for *P. melaninogenica* to enhance *S. pneumoniae* clearance, as 24 hour exposure to other commensal bacteria or *Escherichia coli* lipopolysaccharide (LPS) did not provide the same protective benefit ^26^. Here, we found that neutrophils purified from the lungs of *P. melaninogenica*-exposed mice exhibited an increased capacity to kill *S. pneumoniae* compared to neutrophils from mice treated with *Escherichia coli* LPS (**Fig. 3A**). Neutrophil serine protease activity, a requirement for *S. pneumoniae* killing after *P. melaninogenica* exposure, is similarly low in neutrophils of non-primed and *E. coli* LPS-treated mice ^26^. Therefore, PBS treatment was used as a non-primed, negative control for neutrophil functional assays moving forward. We used an in vivo flow cytometry-based uptake assay, where mice were injected i.t. with pre-opsonized, FITC-labeled HK *S. pneumoniae* 24 hours after PBS or *P. melaninogenica* exposure, to assess the ability of lung neutrophils to phagocytose *S. pneumoniae*. Neutrophil phagocytosis of *S. pneumoniae* was significantly increased in the lungs of *P. melaninogenica*-primed mice (**Fig. 3B**). In addition to phagocytic and killing capacity, neutrophils were analyzed for surface expression of CD18, an integrin subunit which together with CD11b forms complement receptor 3 (CR3), important for phagocytosis of iC3b-opsonized bacteria ^32^. Neutrophil expression of CD18 was significantly higher in *P. melaninogenica*-primed mice (**Fig. 3C**), (**Fig. S5A**), correlating with the increased phagocytosis and *S. pneumoniae* killing. Of note, C3 and other complement associated genes were upregulated in neutrophils and macrophages in *P. melaninogenica*-primed mice (**Fig. S5B**).

**Figure 3:**
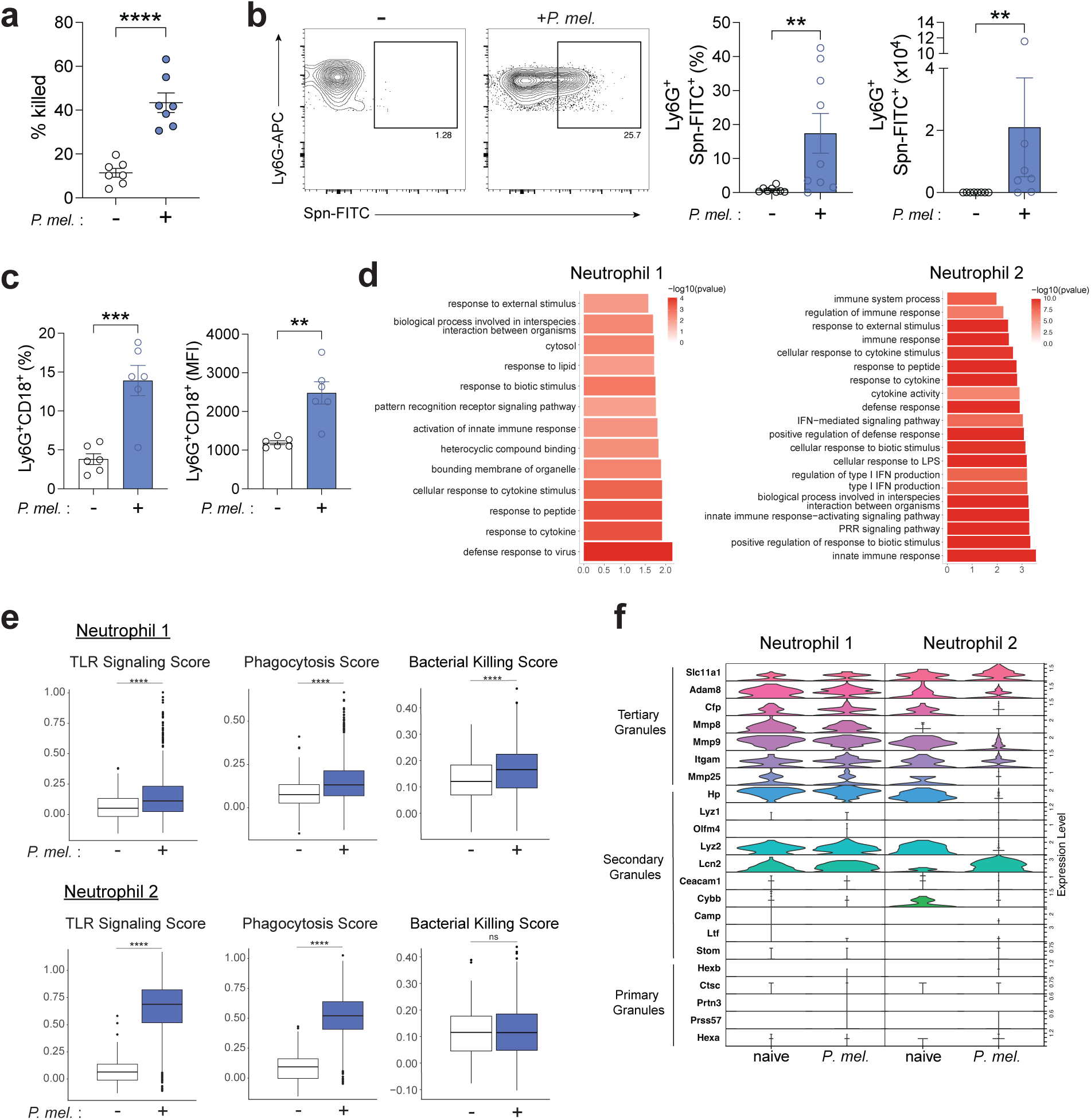
*P. melaninogenica* induces neutrophil transcriptional profiles associated with enhanced phagocytosis and killing of *S. pneumoniae.* **A**) Percent of *Spn* killed by neutrophils isolated from mice exposed to either *E. coli* LPS or *P. mel* HK for 24 hours (mean ± SEM; n = 7 biological replicates pooled from 3 independent experiments). Quantification of Ly6G+Spn-FITC+ (**B**) and Ly6G+CD18+ (**C**) neutrophils in lung tissue of mice exposed to FITC-labeled *Spn* HK for 3 hours, with prior exposure to PBS or *P. mel* HK (mean ± SEM; n = 6-9 biological replicates pooled from 2-3 independent experiments). (**D**) Top differentially enriched GO pathways in neutrophil populations of mice exposed to *P. mel* HK relative to naïve mice. (**E**) Module scoring analysis of toll-like receptor signaling, phagocytosis, and bacterial killing signature genes in neutrophil populations of naïve mice and mice exposed to *P. mel* HK. (**F**) Violin plot displaying expression of primary, secondary, and tertiary granule genes in neutrophil populations of naïve mice and mice exposed to *P. mel* HK. Data were analyzed by unpaired t test (**A,C**), Mann-Whitney test (**B**), or Wilcoxon rank-sum test (**E**), \*\**p*<0.01, \*\*\**p*<0.001, \*\*\*\**p*<0.0001.

These findings prompted us to characterize the transcriptional changes in neutrophils from mice primed with *P. melaninogenica*. Unbiased analysis of the two neutrophil populations identified by scRNA-seq revealed significant enrichment for several gene ontology (GO) terms involved in immune defense, including pattern recognition receptor signaling pathway, response to cytokine stimulus, and activation of innate immune response in *P. melaninogenica*-exposed mice (**Fig. 3D**). To investigate differences in phagocytosis and toll-like receptor signaling associated genes, a module scoring function was applied to assess expression of selected gene sets in aggregate per condition. Module scoring analysis identified a significant upregulation of both phagocytosis signature genes and toll-like receptor signaling genes (**Fig. 3E**), which stimulate neutrophil activation, phagocytosis, and reactive oxygen species production ^33^, in *P. melaninogenica*-exposed mice. Module scoring also indicated increased maturation associated genes in the Neutrophil 1 population, which was smaller in total number compared to the largely expanded Neutrophil 2 population, and significantly elevated migration signature genes in both neutrophil populations in *P. melaninogenica*-exposed mice (**Fig. S5C**). Several neutrophil granule genes were reduced in the Neutrophil 2 population of *P. melaninogenica*-primed mice (**Fig. 3F**), which may indicate reduced granule synthesis in more matured neutrophils ^34,35^. Together, these findings suggest that *P. melaninogenica* exposure leads to transcriptional upregulation of antibacterial defense genes in lung neutrophils, corresponding with increased phagocytosis and killing of *S. pneumoniae*.

#### TNF signaling drives enhanced neutrophil-mediated clearance of *S. pneumoniae* following *P. melaninogenica* priming

In bronchoalveolar lavage fluid from mice exposed to HK *P. melaninogenica*, TNF⍺ was the highest measured cytokine, produced as early as 2 hours post-exposure to *P. melaninogenica* (**Fig. 4A**). At the transcriptional level, neutrophils were the primary *Tnf* expressing cells, with expression also noted in monocyte and macrophage populations, and *Tnf* expression was elevated in *P. melaninogenica*-exposed mice (**Fig. 4B-C**). TNF⍺ is recognized by TNF receptor 1 (TNFR1) and TNFR2, which are encoded by the *Tnfrsf1a* and *Tnfrsf1b* genes, respectively. As expected, *Tnfrsf1a* was broadly expressed across different lung cell types, while *Tnfrsf1b* was primarily expressed by myeloid cells and T regulatory cells ^36^ (**Fig. 4D**). Both TNF receptors were highly expressed by neutrophils. Using CellChat analysis to infer cell-cell interactions, neutrophils were predicted to respond to TNF⍺ produced by Cxcl3+ MDMs and from themselves in an autocrine fashion (**Fig. 4E**). By module scoring analysis, both neutrophil populations in *P. melaninogenica*-exposed mice exhibited significant upregulation of TNF signaling pathway associated genes (**Fig. 4F**). Linear correlation demonstrated a significant positive correlation between expression of phagocytosis-related and TNF signaling pathway genes in the Neutrophil 1 population of *P. melaninogenica*-exposed mice (**Fig. 4G**).

**Figure 4:**
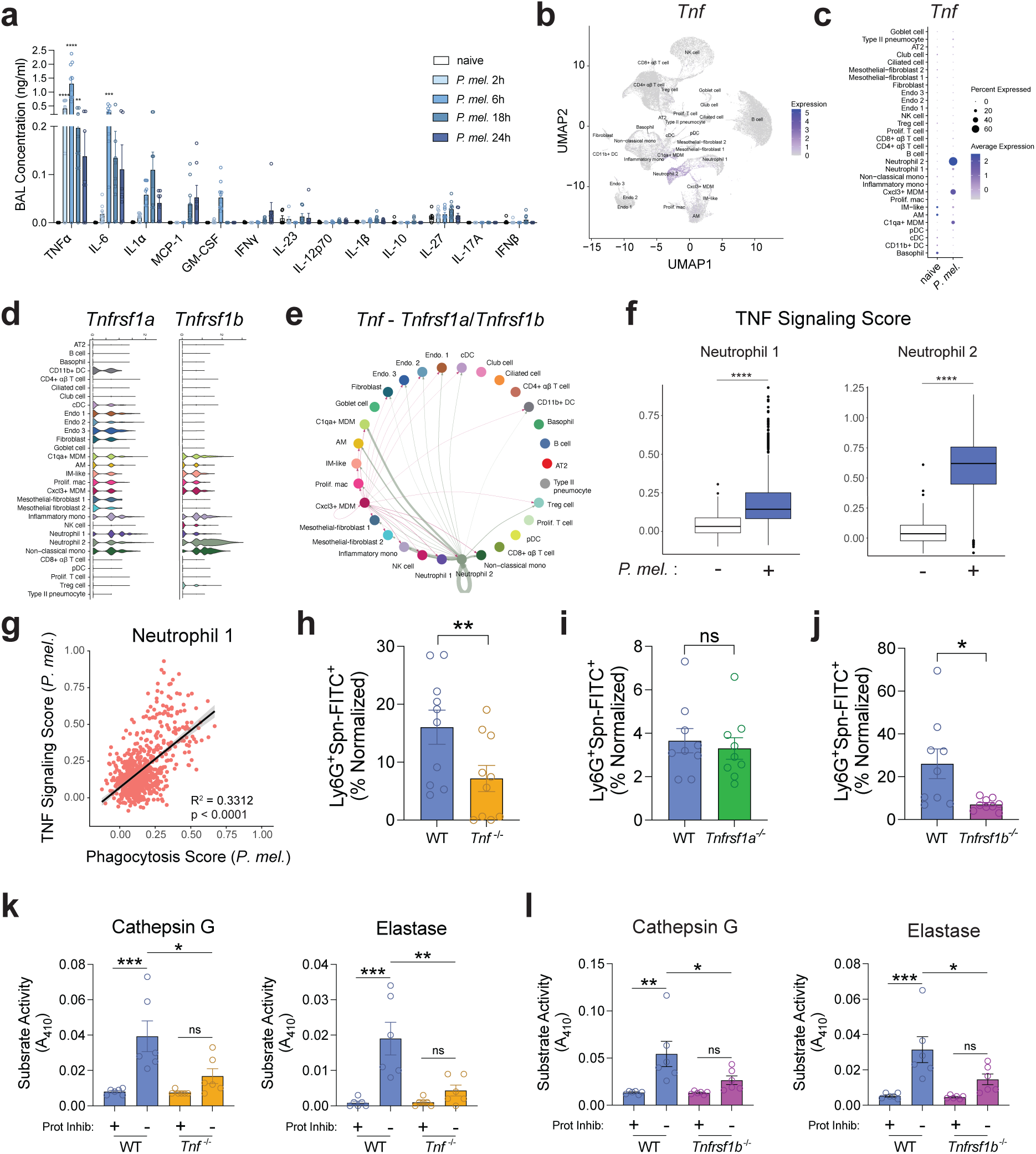
TNF signaling drives enhanced neutrophil-mediated clearance of *S. pneumoniae* following *P. melaninogenica* priming. (**A**) Quantification of cytokines and chemokines in lung BAL of mice exposed to *P. mel* HK for 0, 2, 6, 18, or 24 hours (mean ± SEM; n = 7-9 biological replicates pooled from 3 independent experiments). (**B**) UMAP overlay of *Tnf* gene expression. (**C**) Dot plot showing *Tnf* expression across cell populations in naïve vs *P. mel* HK-exposed mice. (**D**) Violin plot of *Tnfrsf1a* and *Tnfrsf1b* expression across all cell populations. (**E**) CellChat circle plot displaying inferred *Tnf*-*Tnfrsf1a*/*Tnfrsf1b* signaling network. (**F**) Module scoring analysis of TNF signaling signature genes in neutrophil populations of naïve mice and mice exposed to *P. mel* HK. (**G**) Pearson’s correlation of phagocytosis and TNF signaling module scores for Neutrophil 2 population from *P. mel* HK-exposed mice. Quantification of Ly6G+Spn-FITC+ neutrophils isolated from WT vs *Tnf*^-/-^ (**H**) *Tnfrsf1a*^-/-^ (**I**), or *Tnfrsf1b*^-/-^ mice (**J**) and incubated with FITC-labeled *Spn* HK for 30 min (mean ± SEM; n = 9-10 biological replicates pooled from 3-4 independent experiments). Serine protease activity detected in neutrophils purified from WT vs *Tnf*^-/-^ (**K**) or *Tnfrsf1b*^-/-^ (**L**) mice, with or without protease inhibitor cocktail (mean ± SEM; n = 6 biological replicates pooled from 2 independent experiments). Abbreviations: AM = alveolar macrophage, IM = interstitial macrophage, MDM = monocyte-derived macrophage, mono = monocyte, mac = macrophage, DC = dendritic cell, cDC = conventional DC, pDC = plasmacytoid DC. Data were analyzed by two-way ANOVA with Dunnett’s post hoc test (**A**), Mann-Whitney test (**H,I**), unpaired t test (**J**), or ordinary one-way ANOVA with Šídák’s post hoc test (**K,L**). \**p*<0.05, \*\**p*<0.01, \*\*\**p*<0.001, \*\*\*\**p*<0.0001.

Given the correlation between phagocytosis and TNF signaling associated genes, we sought to uncover whether TNF⍺ signaling regulates *S. pneumoniae* phagocytosis in neutrophils. To test this, neutrophils were isolated from the lungs of wildtype and *Tnf*^-/-^mice 24 hours after *P. melaninogenica* exposure and incubated with pre-opsonized, FITC-labeled HK *S. pneumoniae* using an ex vivo uptake assay. In the absence of TNF⍺, neutrophil phagocytosis of *Streptococcus pneumoniae* was significantly reduced (**Fig. 4H**). To determine which TNF⍺ receptors were necessary for enhanced *S. pneumoniae* phagocytosis, uptake of FITC-labeled HK *S. pneumoniae* was evaluated in neutrophils purified from wildtype, *Tnfrsf1a^-/-^*, or *Tnfrsf1b^-/-^* mice following *P. melaninogenica* priming. While phagocytic capacity was retained in neutrophils lacking TNFR1, *S. pneumoniae* phagocytosis was significantly reduced in neutrophils lacking TNFR2 compared to neutrophils from WT mice (**Fig. 4I-J**). Enhanced killing of *S. pneumoniae* after *P. melaninogenica* exposure was previously linked to intracellular serine protease activity ^26^. To extend this finding, we quantified serine protease activity levels in neutrophils isolated from the lungs of wildtype, *Tnf*^-/-^, or *Tnfrsf1b*^-/-^ mice after *P. melaninogenica* priming. Neutrophils lacking either TNF or TNFR2 exhibited significantly reduced serine protease activity (**Fig. 4K-L**). Neutrophil expression of reactive oxygen species, which also contributes to intracellular bacterial killing, was unaffected by the absence of TNF⍺ (**Fig. S5D**). Additionally, loss of TNF⍺ or TNFR2 did not impact neutrophil frequencies in naïve or *P. melaninogenica*-exposed mice (**Fig. S5E-F**). Together, these data identify TNF⍺ signaling through TNFR2 as a previously unrecognized regulator of neutrophil serine protease activity and bacterial uptake in the lung during acute pneumococcal pneumonia.

#### Protective immune priming reduces the pneumococcal-induced type I IFN response

In mice infected with *S. pneumoniae* alone, we identified expression of several genes associated with canonical innate immune activation during pneumococcal infection, including genes involved in interferon signaling, inflammasome signaling, pro-inflammatory cytokine production, and host cell invasion (**Fig. S6A**). Using pathway enrichment analysis as an unbiased approach to identify the top differentially enriched signaling pathways, type I and type II interferon associated pathways emerged as the dominant gene ontology pathways across myeloid populations in *S. pneumoniae*-infected mice (**Fig. 5A**). Interferon-stimulated genes (ISGs) were also among the top upregulated genes in the C1qa+ MDM population which was selectively enriched in *S. pneumoniae* infected mice (**Fig. 5B**) and broadly among myeloid cells (**Fig. 5C**). Correspondingly, expression of type I interferon receptor genes *Ifnar1* and *Ifnar2*, as well as type II interferon receptor gene *Ifngr1*, were higher in myeloid populations compared to other cell types (**Fig. S6B**). Interferon signaling has context-dependent effects during *S. pneumoniae* infection. While interferon signaling can enhance epithelial and innate immune cell defense against *S. pneumoniae* ^37–40^, excessive type I interferon stimulation increases susceptibility to pneumococcal pneumonia through suppression of myeloid cell recruitment and activation ^41–43^. Notably, ISGs were expressed at lower levels in mice exposed to *P. melaninogenica* prior to *S. pneumoniae* infection (**Fig. 5C**), in which enhanced *S. pneumoniae* clearance was observed. While similar interferon-associated genes were upregulated during *S. pneumoniae* infection, particularly among C1qa+ MDMs and neutrophils, module scoring analysis indicated a significant reduction in the expression of both type I and type II interferon signature genes in these cells, as well as inflammatory monocytes and dendritic cells, in mice primed with *P. melaninogenica* prior to *S. pneumoniae* infection (**Fig. 5D**). Thus, *P. melaninogenica* exposure is associated with reduced interferon signaling in myeloid cells during *S. pneumoniae* infection.

**Figure 5:**
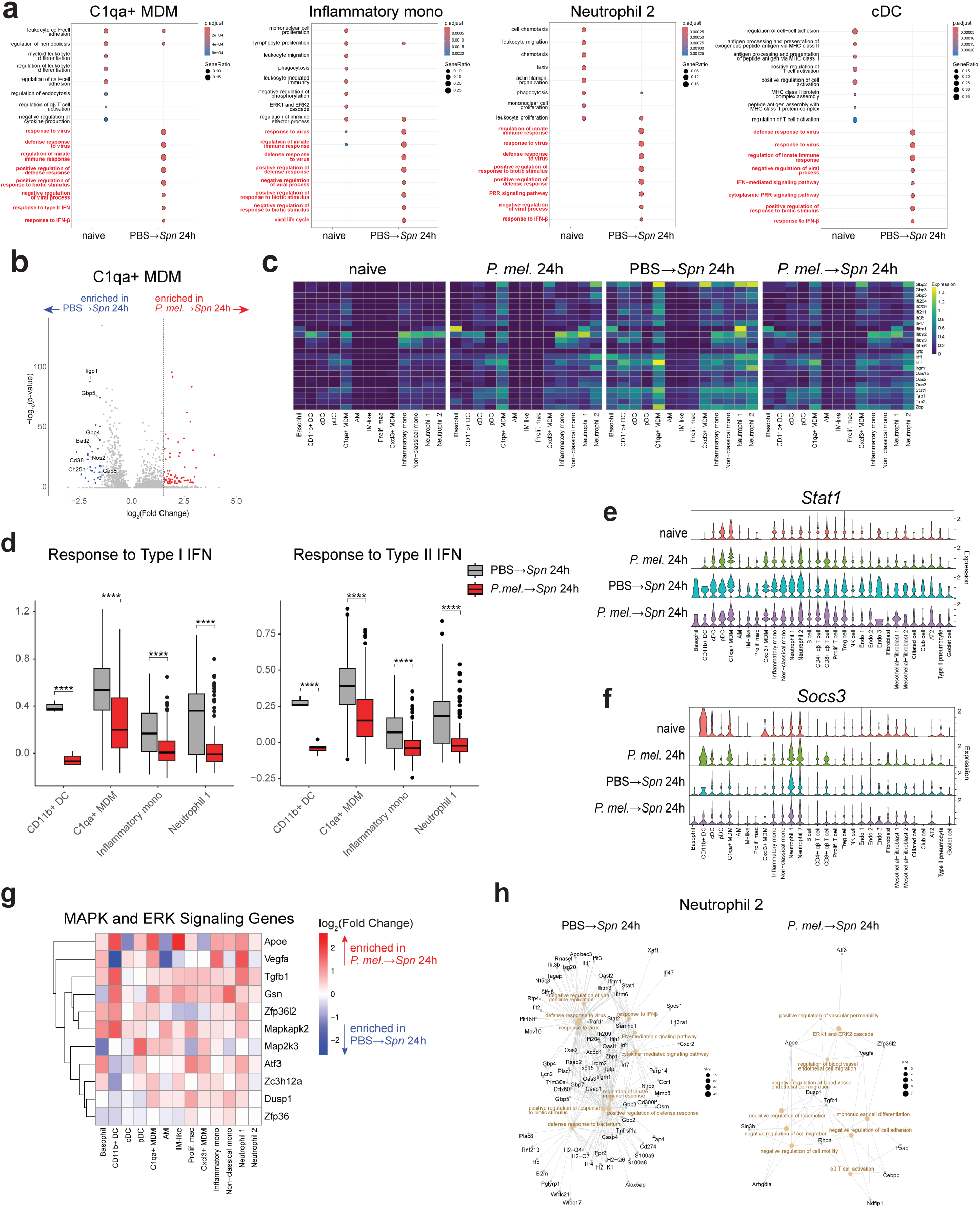
Protective immune priming reduces the pneumococcal-induced IFN response. (**A**) Top enriched GO pathways in selected myeloid cell populations from naïve vs Spn-infected mice. Red text indicates GO Terms in which enriched genes are predominantly ISGs. (**B**) Volcano plot of top enriched genes in C1qa+ MDMs from *Spn*-infected mice with vs without prior *P. mel* HK exposure, with individual ISGs indicated. (**C**) Heatmap of ISG expression across myeloid cell populations, split by condition. (**D**) Module scoring analysis of type I interferon and type II interferon signaling signature genes in myeloid cell populations of *Spn*-infected mice with vs without prior exposure to *P. mel* HK. Violin plots showing expression of (**E**) *Stat1* and (**F**) *Socs3* across all cell types, by condition. (**G**) Heatmap displaying log-fold changes of MAPK signaling gene expression in myeloid cells of mice exposed to *P. mel* HK prior to *Spn* infection, relative to mice infected with *Spn* alone. (**H**) Network plot of top enriched pathways (beige nodes) and associated genes in the Neutrophil 2 population for *Spn*-infected mice with or without prior *P. mel* HK exposure. Data were analyzed by Wilcoxon’s rank-sum test (**D**). \*\*\*\**p*<0.0001.

Expression of *Stat1*, which controls expression of interferon-stimulated genes ^44^, was upregulated in all cell types in the lungs of *S. pneumoniae*-infected mice (**Fig. 5E**). In contrast, suppressor of cytokine signaling 3 (*Socs3*), a negative regulator of the JAK/STAT signaling pathway ^45^, was increased in neutrophils, monocytes, and T cells of *P. melaninogenica*-exposed mice, which could contribute to the regulation of interferon signaling (**Fig. 5F**). Across myeloid cell populations, MAPK and ERK signaling associated genes were upregulated in mice primed with *P. melaninogenica* prior to *S. pneumoniae* infection, compared to mice infected with *S. pneumoniae* alone (**Fig 5G**). Similarly, network pathway analysis revealed ERK1 and ERK2 cascade signaling as a top upregulated pathway in neutrophils from mice primed with *P. melaninogenica* before *S. pneumoniae* infection, compared to several interferon signaling related pathways dominating neutrophils in unprimed mice infected with *S. pneumoniae* (**Fig. 5H**). Of note, MAPK and ERK1/2 signaling occur downstream of TLR2 ^46^, shown to be required for *P. melaninogenica*-mediated protection ^26^. Additionally, genes involved in chemotaxis and phagocytosis, processes known to be regulated by MAPK-ERK signaling ^47–49^, were enriched in *P. melaninogenica*-primed mice (**Fig. S5G**). These data suggest that the improved immune-mediated clearance of *S. pneumoniae* in mice exposed to *P. melaninogenica* involves a skewing of the typical myeloid cell transcriptional response to acute pneumococcal pneumonia away from interferon signaling pathway dominance.

### Monocyte and macrophage populations contribute to *P. melaninogenica*-induced clearance of *S. pneumoniae*

Due to the altered macrophage signaling patterns and distinct MDM populations detected during *S. pneumoniae* infection with versus without *P. melaninogenica* exposure, we next analyzed transcriptional differences in macrophages from primed versus unprimed mice. In C1qa+ MDMs, lysosome and lytic vacuole organization genes involved in phagocytosis and bacterial killing were enriched in mice exposed to *P. melaninogenica* prior to *S. pneumoniae* infection, compared to mice infected with *S. pneumoniae* alone (**Fig. 6A**). Lysosome organization genes were broadly upregulated across all monocyte and macrophage populations in mice exposed to *P. melaninogenica* prior to *S. pneumoniae* infection (**Fig. 6B**). Similarly, unbiased pathway enrichment analysis identified several pathways important for bacterial killing in alveolar macrophages from mice exposed to *P. melaninogenica* before *S. pneumoniae* infection, including lysosomal acidification and reactive oxygen species metabolic process associated genes (**Fig. 6C**). Network pathway analysis confirmed enrichment in lysosome regulation and positive regulation of defense response genes in AMs for *P. melaninogenica*-primed, but not unprimed, mice infected with *S. pneumoniae* (**Fig. 6D**). To determine the functional consequences of these transcriptional differences, we performed an in vivo phagocytosis assay to compare *S. pneumoniae* uptake in mice with or without *P. melaninogenica* exposure. Assessment of phagocytic capacity revealed a significant increase in phagocytosis of FITC-labeled HK *S. pneumoniae* for both recruited Ly6C+ monocytes and MDMs as well as resident alveolar macrophages in mice primed with *P. melaninogenica* (**Fig. 6E**).

**Figure 6:**
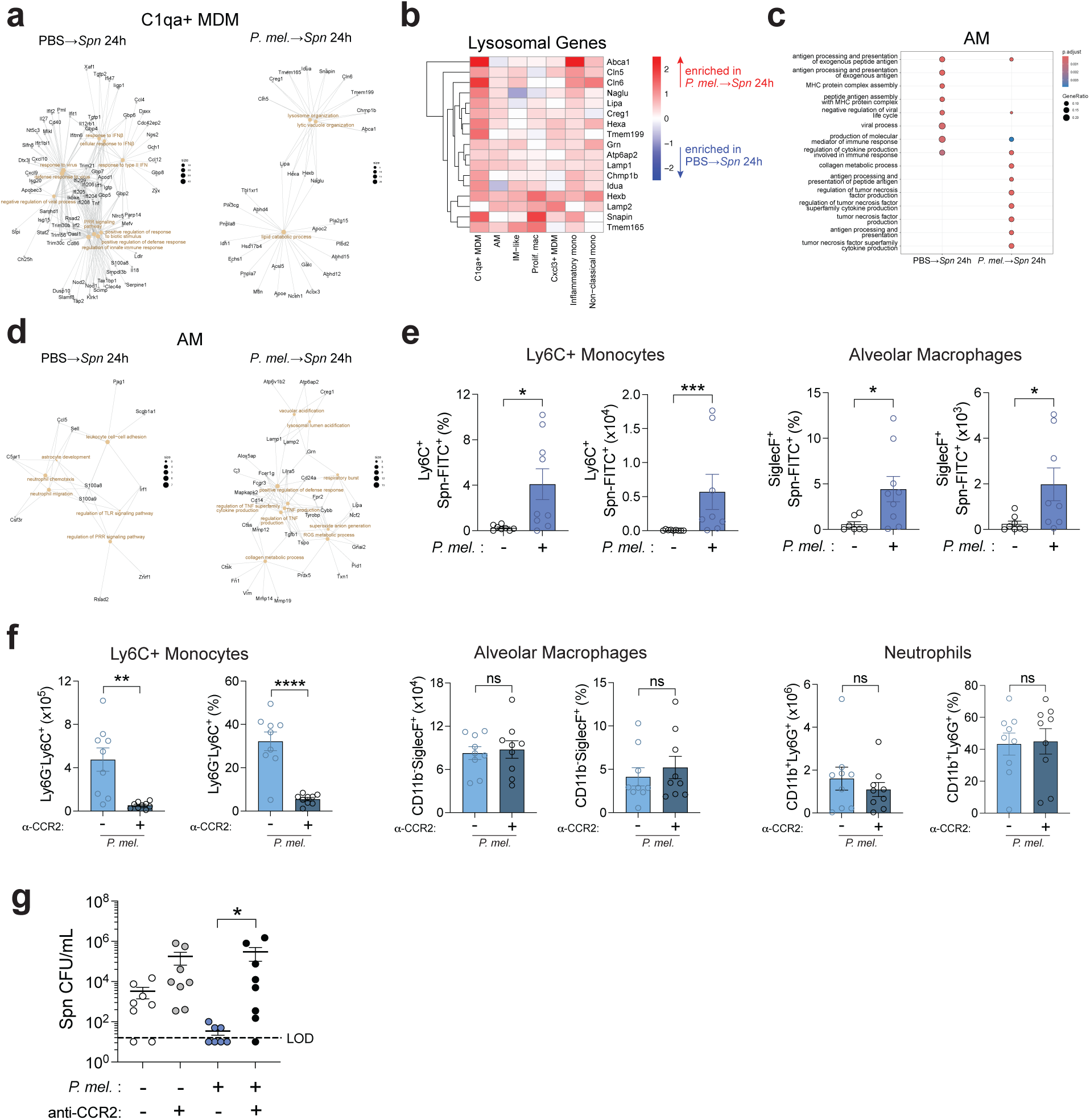
Monocyte and macrophage populations contribute to *P. melaninogenica*-induced clearance of *S. pneumoniae.* (**A**) Network plot of top enriched pathways (beige nodes) and associated genes in C1qa+ MDMs from *Spn*-infected mice with or without prior *P. mel* HK exposure. (**B**) Heatmap of log-fold changes in lysosomal gene expression in monocyte and macrophage populations from mice exposed to *P. mel* HK prior to *Spn* infection, relative to mice infected with *Spn* alone. (**C**) Top enriched GO pathways in AMs from *Spn*-infected mice with or without prior *P. mel* HK exposure. (**D**) Network plot of top enriched pathways and associated genes in AMs from *Spn*-infected mice with or without prior *P. mel* HK exposure. (**E**) Quantification of Ly6C+Spn-FITC+ monocytes and SiglecF+Spn-FITC+ AMs in lung tissue of mice exposed to FITC-labeled *Spn* HK for 3 hours, with or without prior exposure to *P. mel* HK (mean ± SEM; n = 7-9 biological replicates pooled from 3 independent experiments). (F) Quantification of Ly6G-Ly6C+ monocytes, CD11b-SiglecF+ AMs, and CD11b+Ly6G+ neutrophils in mice treated with isotype control or anti-CCR2 antibodies prior to *P. mel* HK exposure (mean ± SEM; n = 9 biological replicates pooled from 3 independent experiments) (**G**) Lung Spn burdens in mice treated with isotype control or anti-CCR2 antibody (MC-21) i.p. together with PBS or *P. mel* HK exposure prior to *Spn* infection (1×10^7^ CFU) (mean ± SEM; n = 8-9 biological replicates pooled from 3 independent experiments). Data were analyzed by Mann-Whitney test (**E,F**), unpaired t test (**E,F**), or Kruskal-Wallis with Dunn’s post hoc test (**G**). \**p*<0.05, \*\**p*<0.01, \*\*\**p*<0.001.

Given the increased bacterial uptake, we sought to understand whether recruited monocytes and macrophages directly contribute to enhanced clearance of *S. pneumoniae* in *P. melaninogenica*-primed mice. To address this question, mice were treated with an anti-CCR2 antibody, which selectively depleted recruited monocytes and macrophages without affecting resident AMs or recruited neutrophil populations (**Fig. 6F**). In the absence of recruited monocytes and macrophages, *P. melaninogenica* exposure was no longer protective against *S. pneumoniae* infection (**Fig. 6G**). Notably, neutrophil activation, as measured by CD18 surface expression and intracellular TNF⍺, was not affected by anti-CCR2 treatment (**Fig. S7A-B**), suggesting that regulation of neutrophil-mediated bacterial killing was not the mechanism responsible for CCR2-dependent enhanced protection against *S. pneumoniae* infection in *P. melaninogenica*-primed mice. Together, these data reveal a previously unrecognized dependence on recruited monocytes and macrophages for commensal-enhanced protection against pneumococcal pneumonia.

## Discussion

Using a preclinical model recapitulating associations between *Prevotella* aspiration and improved outcomes in critically ill patients, we demonstrate that *P. melaninogenica* dramatically alters the pulmonary transcriptional innate immune response to *S. pneumoniae*. Differential recruitment and activation of monocyte-derived macrophage populations with opposing antibacterial signaling programs was linked to improved pathogen clearance following *Prevotella* exposure. Further, we discovered an essential role for TNFR2 in neutrophil-mediated protection during bacterial pneumonia, mediated by increased neutrophil phagocytosis and serine protease activity. Together, these findings illustrate a new paradigm for improved antibacterial defense through commensal-directed myeloid cell reprogramming. In prior work, we defined the unique microbial requirements for *P. melaninogenica* enhancement of the acute immune response to *S. pneumoniae* lung infection^26^. While live or HK *P. melaninogenica* and several other *Prevotella* species demonstrated similar enhanced protection against *S. pneumoniae* infection, another species, *Prevotella intermedia*, was not protective and failed to induce a similar TLR2-dependent neutrophil TNFα response as the ‘protective’ species^26^. Moreover, neither HK preparations of other airway commensal bacteria nor *E. coli* LPS were protective, indicating a specificity beyond Gram-negative LPS or bacterial exposure. It isn’t yet clear how broadly *P. melaninogenica* protection extends, though we have shown enhanced protection against both serotype 2 and serotype 3 *S. pneumoniae* as well as another clinically relevant Gram-positive pathogen, *Staphylococcus aureus*^26,27^. For this study, we focused on defining the transcriptional reprogramming induced by exposure to HK *P. melaninogenica* as an extremely common microbial aspiration which may precede *S. pneumoniae* infection. The enhanced immune response to pneumococcal infection observed following *P. melaninogenica* exposure also provided the opportunity to uncover the transcriptional changes associated with improved *S. pneumoniae* clearance, compared to less robust infection defense.

To our surprise, the dominant transcriptional signature enriched in myeloid cells was an anti-viral associated interferon response. *S. pneumoniae* triggers type I interferon signaling via the pore-forming toxin pneumolysin ^37^, which perforates host cell membranes leading to detection of bacterial DNA in the cytosol through the c-GAS STING pathway ^50^. While interferon signaling can protect against *S. pneumoniae* infection by increasing alveolar epithelial cell survival and myeloid cell activation ^38,39^, excessive type I interferon signaling is widely documented to increase susceptibility to pneumococcal pneumonia and worsen pneumonia outcomes, particularly in the context of viral co-infections ^41–43,51^. In *P. melaninogenica*-primed mice, the interferon response was significantly subdued compared to mice infected with *S. pneumoniae* alone. In this setting, expression of the transcription factor *Socs3*, which inhibits JAK/STAT signaling, was elevated in C1qa+ MDMs and neutrophils, the two cell types exhibiting the strongest ISG response following *S. pneumoniae* infection. These data suggest that *Socs3* expression, specifically within ISG-expressing myeloid cells, may regulate the otherwise dominant interferon response. The effect of type I interferon on phagocyte activation is variable, as vector delivery of IFNα was shown to improve *S. pneumoniae* killing while others studies indicate inhibitory effects ^39,42,43^. In myeloid cells from mice primed with *P. melaninogenica* prior to infection, we observed overall reduced ISG expression and a corresponding increase in the expression of MAPK and ERK signaling genes, aligning functionally with increased *S. pneumoniae* uptake by lung neutrophils and macrophages. Phagocytosis of *S. pneumoniae* by innate immune cells is a critical step for bacterial killing, as the primary effective killing mechanisms occur intracellularly^52^. Together, these data suggest that *P. melaninogenica* priming shifts the prototypical *S. pneumoniae*-induced inflammatory response towards a TLR2-driven MAPK and ERK signaling program that supports improved antibacterial defense.

The recruitment of monocyte-derived macrophages to the lungs is a hallmark of severe pneumonia, with direct contributions to alveolar injury ^53–55^. We observed striking differences among the two transcriptionally distinct MDM populations identified by scRNA-seq between mice infected with *S. pneumoniae* compared to those primed with *P. melaninogenica*. The C1qa+ MDM population which dominated *S. pneumoniae*-infected mice were less mature than the Cxcl3+ MDMs and exhibited significantly higher expression of ISGs. C1qa+ MDMs may therefore serve as a key source of the maladaptive interferon response observed in susceptible mice. In people with severe bacterial pneumonia, recent reports indicate a central role for circulating C1qa-expressing monocytes in driving deleterious inflammation ^56,57^. In contrast, the Cxcl3+ MDM population selectively enriched by *P. melaninogenica* exposure transcriptionally resembled mature tissue resident-like macrophages. The importance of monocytes and MDMs for *P. melaninogenica*-mediated protection was confirmed by CCR2-depletion, which abrogated the protective effect of *Prevotella* immune priming. While CCR2-depletion cannot distinguish between Cxcl3+ and C1qa+ MDMs, their transcriptional differences indicate the potential for unique roles during infection defense, an important area for further study. Other examples of enhanced defense against bacterial pneumonia following immune priming by microbial-derived factors, gut commensals, or other pneumococcal serotypes are all mechanistically linked to reprogramming in resident macrophages, namely AMs ^58–62^. Here, our findings highlight an alternative mechanism for enhanced defense against bacterial pneumonia, through altered MDM recruitment and activation during acute infection.

In addition to macrophages, we identified two neutrophil populations by scRNA-seq with distinct responses to *P. melaninogenica* exposure. Neutrophils undergo several maturation states which change following tissue migration and during bacterial infection ^63^, with two dominant populations reported during murine *S. pneumoniae* infection and for infected human lungs ^64,65^, similar to our scRNA-seq dataset. Of the two populations identified, Neutrophil 2 expressed higher levels of inflammatory genes, including those involved in TLR signaling, phagocytosis, and TNF signaling, in response to *P. melaninogenica*. At the functional level, neutrophil phagocytosis and surface activation marker expression increased following *P. melaninogenica* airway exposure; however, neutrophil subpopulations could not be purified or distinguished from one another based on surface protein alone. Notably, neutrophil activation as measured by CD18 expression and TNF⍺ production did not change in the absence of recruited monocytes. These data suggest that either signaling from non-monocyte populations regulate neutrophil activation after *P. melaninogenica* exposure, or neutrophils activate themselves in an autocrine fashion. While TNFR1 is activated by soluble TNF, TNFR2 is activated primarily by membrane-bound TNF (mTNF) on the surface of adjacent cells ^66^. Binding of mTNF to TNFR2 induces reverse signaling through the ligand (mTNF)-bearing cell ^67^. This may explain why isolated neutrophils lacking TNFR2 exhibited reduced bacterial uptake even though they were able to produce TNF⍺. mTNF signaling through TNFR2 is a pro-survival signal in neutrophils, in contrast to TNFR1, which promotes apoptosis ^68^. However, the independent contributions of TNFR1 versus TNFR2 signaling during infection are largely unclear. During skin infection with *Staphylococcus aureus*, TNFR2 was protective through activation of neutrophil ROS and vital NETosis ^69^. Similarly, we identified a non-redundant role for TNFR2 in neutrophil-mediated protection during bacterial pneumonia, though in this case through increased neutrophil phagocytosis and serine protease activity without altered ROS. Previously, we found that TNF⍺ blockade abrogated *P. melaninogenica* enhanced protection against *S. pneumoniae* lung infection^26^, indicating the importance of TNF signaling for pathogen clearance. While *P. melaninogenica* exposure can directly induce neutrophil production of TNF⍺ ex vivo, additional tissue-specific cells or signals are required for enhanced *S. pneumoniae* killing^26^. Here, we cannot rule out the contribution of TNFR2 expression on other cells for enhanced neutrophil antimicrobial function. Together, these findings highlight TNF signaling through TNFR2 as a critical regulator of neutrophil serine protease activity, which is a key mechanism of *S. pneumoniae* killing^70,71^.

The concept of “trained immunity” describes a functional reprogramming of immune cells and is increasingly recognized as a mechanism which can improve baseline defense against pathogens. During trained immunity, a primary inflammatory stimulus induces epigenetic and metabolic reprogramming in innate immune cells that reshapes subsequent inflammatory responses ^72^. While the long-term consequences of the differentially enriched MDMs observed in resistant versus susceptible groups are unknown, the transcriptional programming in recruited MDMs was associated with a significant impact on bacterial clearance during acute infection, with recruitment of more ‘optimal’ MDMs associated with better infection defense. Although neutrophils are short-lived, emerging work has shown that epigenetic remodeling can occur in bone marrow progenitors after administration of β-glucan or the BCG vaccine in mice, leading to central trained immunity in neutrophils ^73–75^. In our dataset, expression of genes linked to immune training in myeloid cell populations of *P. melaninogenica*-exposed mice were increased, including lipid metabolism genes in macrophages and TLR signaling in both macrophages and neutrophils. Because downstream epigenetic and metabolic adaptations require longer than 24 hours to take effect, extending the time between *P. melaninogenica* exposure and *S. pneumoniae* infection will be necessary to better understand whether trained immunity contributes to enhanced pathogen clearance in this setting. Regardless, our findings identify *P. melaninogenica* exposure as a previously unrecognized microbial stimulus capable of broadly reprogramming myeloid cell responses to bacterial infection.

This study has several limitations to consider. Many immune defense mechanisms are regulated post-transcriptionally, including neutrophil protease secretion and toll-like receptor signaling; therefore, gene expression data were complemented with protein expression analysis and functional assays whenever possible. The *P. melaninogenica* exposure model used here is a single-dose exposure to facilitate characterization of the initial response pathways involved in enhanced host defense against pneumonia. While single pre-exposure models of inflammatory stimuli are commonly used to delineate immune priming mechanisms in murine models^25,74,76,77^, future work could expand these findings by administration of multiple smaller doses over an extended period prior to *S. pneumoniae* infection, mimicking the airway environment where the lungs are continuously exposed to aspirated oral commensals. Another potential limitation is the use of heat-killed *P. melaninogenica* rather than live bacteria. Recent work demonstrated that enrichment of oral commensals in the lower airway correlates with the detection of immune-modulating short-chain fatty acids ^78^. Using a multi-omics approach, the same group showed that *P. melaninogenica* is metabolically active in the lung and produces metabolites that contribute to immune modulation including adenosine, inosine, methionine, and glutamate ^78^. Their findings provide initial evidence that aspirated oral commensals are not merely transient passengers but active participants shaping the lung immune environment. Although we observe enhanced *S. pneumoniae* clearance after priming with either live or heat-killed *P. melaninogenica* ^26^, the full spectrum of immunomodulatory properties of metabolically active *P. melaninogenica* may not be fully recapitulated using heat-killed bacteria.

Overall, these findings highlight key transcriptional innate immune signaling pathways associated with improved antibacterial defense following exposure to a commonly aspirated airway commensal through the enrichment of alternatively activated MDMs, licensing neutrophil serine protease activity via TNFR2, and shifting from an IFN-dominant response to a pro-phagocytic transcriptional program, together correlating with improved *S. pneumoniae* uptake and clearance. Clinically, escalating evidence suggests that airway anaerobes including *Prevotella* should be spared during treatment of critically ill patients, based on the strong associations between anaerobes and improved survival ^14,79–81^. While arguments for this approach are often centered on the potential contributions of gut anaerobes, a recent study concluded that oral and lung anaerobes, rather than gut anaerobes, are the most powerful microbial indicator of improved survival in critically ill patients ^82^. In the same study, *Prevotella* inversely corelated with acute respiratory failure and markers of tissue injury including soluble TNFR1 ^82^. This observation is in alignment with the protective TNF signaling through TNFR2, rather than pro-apoptotic TNFR1, observed in *P. melaninogenica* exposed mice. Our data, together with the clinical correlation between *P. melaninogenica* abundance and reduced lower airway infections ^13,16,21^, point to an important role for *P. melaninogenica* aspiration in directing lung immune homeostasis and defense against bacterial pneumonia. Further work is needed to optimize *Prevotella*-mediated enhanced defense in a clinical setting, with the potential to reduce overall pneumonia mortality.

## Methods

### Animals

Adult male and female mice aged 7-9 weeks were used for this study. C57BL/6 J (WT), B6.129S-Tnf ^tm1Gkl^/J (*Tnf*^-/-^), C57BL/6-Tnfrsf1a^tm1lmx^/J (*Tnfrsf1a*^-/-^), and B6.129S2-Tnfrsf1b^tm1Mwm^/J (*Tnfrsf1b*^-/-^) mice were purchased from The Jackson Laboratory (strains #000664, #005540, #003242, and #002620, respectively). All mouse strains used in these studies (WT, *Tnf*^−/−^, *Tnfrsf1a*^−/−^, and *Tnfrsf1b*^-/-^) are on the C57BL/6J genetic background. Mice were maintained in the University of Colorado Office of Laboratory Animal Resources.

### Bacteria

*Prevotella melaninogenica* strain ATCC^®^ 25845 was purchased from the American Type Culture Collection (Manassas, VA). To prepare heat-killed *Prevotella* stocks, *P. melaninogenica* was grown on Brucella agar plates anaerobically for 2 days, then resuspended in PBS and incubated at 65°C for 30 min. A streptomycin resistant variant of serotype 2 *Streptococcus pneumoniae* strain D39 was used for these studies^83^. *S. pneumoniae* was grown statically in Todd Hewitt Broth with 5% Yeast Extract (BD Bacto™), with 50 μg/mL streptomycin (Sigma), at 37 °C with 5% CO_2_. Heat-killed *S. pneumoniae* was prepared by growing to OD_600_ > 0.5, resuspending in PBS, and incubating at 65°C for 1 hour. Samples before and after heat-killing were used to determine CFU equivalents/mL and confirm killing, respectively. To prepare FITC- labeled *S. pneumoniae*, heat-killed bacteria were pelleted and resuspended in a 0.5mg/ml FITC solution in PBS and incubated for 1 hour at 4°C. HK *S. pneumoniae* was washed twice with Hank’s Balanced Salt Solution (Gibco) + 1% bovine serum albumin (Fisher BioReagents™) and then resuspended in HBSS + 1% BSA.

### Infections

For *S. pneumoniae* infections, bacteria were grown to mid-log phase in Todd Hewitt Broth with 5% Yeast Extract, centrifuged at ≥20,000 x *g* for 10 min, and resuspended in PBS before storing at -80°C. CFU plates were used to determine concentration of frozen stocks. For infections, *S. pneumoniae* frozen stocks were thawed and resuspended at desired concentration, and mice were administered 10^7^ CFU *S. pneumoniae* i.t. in 50 μL PBS. For *P. melaninogenica* exposures, mice were administered 50 μL i.t. of 10^6^-10^8^ CFU in PBS 24 hours prior to *S. pneumoniae* infection. For CFU enumeration, lung tissue was homogenized using the Bullet Blender tissue homogenizer (Stellar Scientific). Lung homogenates were serially diluted and plated on blood agar plates (Remel). For CCR2+ monocyte depletion, mice were intraperitoneally injected with either 20 μg MC-21 antibody ^84^ or rat IgG2B isotype control (Bio X Cell; catalog #BE0090, clone LTF-2, lot #915224J3) resuspended in 200 μL PBS 24 hours prior and immediately before treating with HK *P. melaninogenica* i.t.

### Flow Cytometry

Lungs were harvested following perfusion by transcardial injection of 10 mL PBS, and single cells were prepared for flow cytometry ^26^. Lung tissue was minced and then incubated in enzymatic digestion buffer (DNaseI 30 μg/mL, Sigma, and type 4 collagenase 1 mg/mL, Worthington Biochemical Corporation, in Hank’s Balanced Salt Solution with calcium and magnesium) for 20 min at 37°C with 5% CO_2_. Tissue was strained through a 70 μM cell strainer (Falcon). To remove red blood cells, samples were pelleted and resuspended in 1mL RBC lysis buffer (0.15 M NH_4_Cl, 10 mM KHC0_3_, 0.1 mM Na_2_EDTA, pH 7.4) for 2 min. Fc receptors were blocked by incubation in anti-CD16/32 (BioLegend) prior to staining in FACS buffer (1% BSA, 0.01% NaN3, PBS). For reactive oxygen species experiments, cells were incubated with Dihydrorhodamine 123 (Sigma) for 20 min at 37°C prior to staining. For intracellular staining, samples were incubated with Brefeldin A (BD Biosciences) diluted 1:1000 for 3 hours at 37°C with 5% CO_2_. Samples were permeabilized with 1 mg/mL saponin (Sigma) in FACS buffer prior to intracellular staining. Antibodies used for staining included anti-mouse: CD45.2 (BioLegend, catalog #564616, clone 104, lot #4339788), CD18 (BD, catalog #749471, clone M1 8/2, lot #5084986), Siglec F (BD, catalog #562681, clone E50-2440, lot #0269846), Ly6C (BioLegend, catalog #128033, clone HK1.4, lot #B432613), Ly6G (BioLegend, catalog #127614, clone 1A8, lot #B366717), CD11b (BioLegend, catalog #101226, clone M1/70, lot #B445235), and TNFα (Invitrogen, catalog #25-7321-82, clone MP6-XT22, lot #2250799). All cells were fixed in 1% paraformaldehyde for 20 min at room temperature. Flow cytometry was performed on the LSR Fortessa X-20 and the Cytek Aurora in the Cancer Center Flow Cytometry Shared Resource Laboratory at the University of Colorado Anschutz Medical Campus (RRID: SCR_022035). Analysis of flow cytometry data was performed using FlowJo™ Software, version 10.10.0 (BD Life Sciences).

### Cytokine and chemokine analysis

BAL cytokines and chemokines except for CXCL1, CXCL2, and CXCL5 were quantified via the LEGENDplex™ Mouse Inflammation Panel (BioLegend) and the LEGENDplex™ Mouse Chemokine Panel (BioLegend). Analytes were measured on the LSR Fortessa X-20 in the Cancer Center Flow Cytometry Shared Resource Laboratory at the University of Colorado Anschutz Medical Campus (RRID: SCR_022035). Data were analyzed using the LEGENDplex™ Data Analysis Software Suite (BioLegend). BAL CXCL1, CXCL2, and CXCL5 were measured using the Mouse CXCL1/KC, Mouse CXCL2/MIP-2, and Mouse CXCL5/LIX DuoSet ELISA kits (R&D Systems), respectively. Analytes were measured on a Synergy™ HT Microplate Reader (BioTek), and data were analyzed via GraphPad Prism.

### Neutrophil Functional Assays

For ex vivo opsonophagocytic assays, neutrophils were isolated from the lungs of WT, *Tnf*^-/-^, *Tnfrsf1a*^-/-^, or *Tnfrsf1b*^-/-^ mice following 24hr treatment with HK *P. melaninogenica* i.t., as indicated. Ly6G+ neutrophils were isolated via magnetic bead positive selection (MojoSort PE positive selection kit, BioLegend). In prior work, similar results were observed using positive or negative selection to isolate lung neutrophils ^26^. For opsonization, 2×10^6^ CFU of FITC-labeled HK *S. pneumoniae* were incubated with 7.6% baby rabbit serum (MP Biomedicals) for 30 min at 37°C. HK *S. pneumoniae* was then incubated with 10^5^ neutrophils in HBSS + 1% BSA for 30 min at 37°C while rotating, prior to flow cytometry staining. To measure phagocytosis, FITC+ neutrophils were quantified via flow cytometry using the LSR Fortessa X-20 in the Cancer Center Flow Cytometry Shared Resource Laboratory at the University of Colorado Anschutz Medical Campus (RRID: SCR_022035). For serine protease assays, neutrophils were isolated via positive selection from the lungs of WT, *Tnf*^-/-^, or *Tnfrsf1b*^-/-^ mice following 24-hour treatment with HK *P. melaninogenica* i.t., as indicated. 10^5^ purified neutrophils were incubated with or without 1x HALT™ protease inhibitor cocktail (Thermo Fisher) for 30 min at 37°C while rotating. Cells were washed and lysed with 0.1% Triton X-100. Elastase substrate (0.85 mM MeOSuc-Ala-Ala-Pro-Val-*p*NA, Sigma) and cathepsin G substrate (0.1 mM Succinyl-Ala-Ala-Pro-Phe-pNA, Sigma) were added to neutrophil lysates and incubated for 45 min at 37°C in the dark. Substrate activity was measured by reading absorbance at 410nm on the Synergy™ HT Microplate Reader (BioTek) minus control wells with no neutrophils added. Data was analyzed via GraphPad Prism.

### Fixed Single-cell RNA Sequencing

Mouse lung tissue was minced and incubated in digestion buffer (DNaseI 0.4 mg/mL, Sigma, type 4 collagenase 1.5 mg/mL, Worthington Biochemical Corporation, 5% fetal bovine serum, CPS Serum, and 10mM HEPES Buffer, Gibco, in Hank’s Balanced Salt Solution with calcium and magnesium) for 20 min at 37°C with 5% CO_2_. Samples were then processed as described in the Flow Cytometry section. Cells with greater than 80% viability were fixed in a 4% formaldehyde fixative solution using the 10X Genomics Chromium Next GEM Single Cell Fixed RNA Sample Preparation Kit. The whole transcriptome probe pairs (10X Genomics) were added to the fixed single cell suspensions to hybridize to their complementary target RNA in an overnight incubation at 42°C. After hybridization, unbound probes were removed with a washing step. The fixed and probe-hybridized single cell suspensions were then loaded onto the Chromium X (10X Genomics) to generate partitioned nanoliter-scale droplets in oil emulsion, with each droplet containing a barcoded bead gel, a single cell, and enzyme Master Mix (10X Genomics) for probe pair ligation and gel bead primer barcode extension. The droplets in oil emulsion were placed in a thermal cycler for 60 min at 25°C, 45 min at 60°C, and 20 min at 80°C. The single cell-barcoded, ligated probe products underwent library preparation using standard 10X Genomics protocols to ensure compatibility with Illumina next-generation sequencing. The gene expression library derived from single cell-barcoded, ligated probe products were sequenced as paired-end 150-base pair reads on the Illumina NOVASeq6000 (Illumina) at the University of Colorado Genomics Shared Resource core, Aurora, CO, USA (RRID: SCR_021984) at a depth of 20,000 reads per cell (1.34B reads total).

### Data Analysis

FASTQ files were processed using CellRanger *multi* along with the Chromium Mouse Transcriptome Probe Set (v1.0.1 mm10-2020-A). Per-sample output feature matrices were used as the input to Seurat (v5) for further analysis. Count matrices were exported for analysis by Scrublet^85^ to infer potential doublets. Scrublet output were added as metadata to the Seurat object. Low quality and potential doublet cells were filtered using parameters: nCount RNA > 500 or <10000, nFeature_RNA >250, percent.mt < 15, doublet score <= 90^th^ percentile. Dataset normalization and scaling was performed using ‘SCTransform’ and was visualized by PCA and UMAP (using 30 dimensions). Integration across conditions was performed using the RPCA method. Nearest-neighbor graph was constructed using 30 dimensions, the integrated reduction, and *k* =30. Initial clustering was performed using a resolution of 0.2. Based on cluster markers, cells were classified as myeloid, stromal, endothelial, epithelial, or lymphocyte. Each subset was then further subclustered at a resolution of 0.1 (except myeloid lymphocytes at 0.2). Cell identities were assigned using a combination of module scoring with known canonical cell type markers, cell type annotation tools (Annotation of Cell Types ^86^; input as top 30 cluster markers per cluster), and manual identification.

Trajectory inference analysis of the macrophage subset was performed using PAGA ^87^. An AnnData object was created from the Seurat object while retaining cell identities, PCs, and variable features. Gene set enrichment analysis and over representation analysis was performed using the GO:ALL database of ClusterProfiler ^88^ and significant cluster markers or DEGs between conditions as input. Cell communication pathways were inferred using CellChat v2 ^89^. Module scoring for specific pathways was calculated using the ‘AddModuleScore’ function of Seurat. Custom gene lists used for modules are included in Source Data. For the Neutrophil module correlation, module scores were plotted and fit with a linear model. For all analyses, default parameters were used unless otherwise specified.

### Study Approval

These studies were approved by the Animal Care and Use Committee of the University of Colorado School of Medicine (protocol #00927) and by the Institutional Biosafety Committee (protocol #1418).

### Statistical Analysis

For biological experiments, statistical analyses were performed using GraphPad Prism. Significant outliers were detected using the ROUT method (Q = 1%) and removed. All data were tested for normality via the Shapiro-Wilk test. For normally distributed data, unpaired t test, one-way ANOVA with Dunnett’s or Šídák’s post-hoc analyses, or two-way ANOVA with Dunnett’s post-hoc analysis were used as specified. For non-Gaussian distributed data, Mann-Whitney tests or Kruskal-Wallis tests with Dunn’s post-hoc analysis were used. For all statistical tests, *p*-values < 0.05 were considered significant.

## Data availability

Transcriptomic data can be accessed in the NCBI Gene Expression Omnibus (GEO) under accession number GSE313404. All other source data are available in the main text and supplementary materials.

## Acknowledgments

The authors would like to acknowledge valuable input on the development of this project from their colleagues at the University of Colorado. These studies were funded by the National Institute of Allergy and Infectious Diseases grant R01AI172958 (SEC), the National Institute on Deafness and Other Communication Disorders grant T32DC012280 (SNS), and the National Institute of Allergy and Infectious Diseases grant T32AI007405 (SNS).

## Author contributions

S.N.S. performed experiments, contributed to experimental design, methodology, data analysis and validation, and wrote the original draft. S.F., S.C.S., and E.R.F. performed experiments. E.L. performed scRNA-seq bioinformatics analysis. E.N.J. assisted with data analysis and methodology. M.M. contributed reagents and methodology. S.E.C. conceived the project, contributed to experimental design, and supervised the project.

## References

1. Bjarnason, A. et al. Incidence, Etiology, and Outcomes of Community-Acquired Pneumonia: A Population-Based Study. Open Forum Infect Dis 5, ofy010 (2018).

2. Hansen, K. et al. Exploring the microbial landscape: uncovering the pathogens associated with community-acquired pneumonia in hospitalized patients. Front Public Health 11, 1258981 (2023).

3. Nicholson, L. K. et al. Clinical and Microbial Determinants of Upper Respiratory Colonization With Streptococcus pneumoniae and Native Microbiota in People With Human Immunodeficiency Virus Type 1 and Control Adults. J Infect Dis 230, 1456–1465 (2024).

4. Yahiaoui, R. Y. et al. Prevalence and antibiotic resistance of commensal Streptococcus pneumoniae in nine European countries. Future Microbiol 11, 737–744 (2016).

5. Abdullahi, O. et al. The prevalence and risk factors for pneumococcal colonization of the nasopharynx among children in Kilifi District, Kenya. PLoS One 7, e30787 (2012).

6. Wu, B. G. & Segal, L. N. The Lung Microbiome and Its Role in Pneumonia. Clin Chest Med 39, 677–689 (2018).

7. Kitsios, G. D. et al. Respiratory Tract Dysbiosis Is Associated with Worse Outcomes in Mechanically Ventilated Patients. Am J Respir Crit Care Med 202, 1666–1677 (2020).

8. Pérez-Cobas, A. E., Ginevra, C., Rusniok, C., Jarraud, S. & Buchrieser, C. The respiratory tract microbiome, the pathogen load, and clinical interventions define severity of bacterial pneumonia. Cell Rep Med 4, 101167 (2023).

9. Dickson, R. P. et al. Bacterial Topography of the Healthy Human Lower Respiratory Tract. mBio 8, e02287–16 (2017).

10. Bassis, C. M. et al. Analysis of the upper respiratory tract microbiotas as the source of the lung and gastric microbiotas in healthy individuals. mBio 6, e00037 (2015).

11. Kitsios, G. D. et al. The upper and lower respiratory tract microbiome in severe aspiration pneumonia. iScience 26, 106832 (2023).

12. Sakwinska, O. et al. Nasopharyngeal microbiota in healthy children and pneumonia patients. J Clin Microbiol 52, 1590–1594 (2014).

13. de Steenhuijsen Piters, W. A. A., et al. Dysbiosis of upper respiratory tract microbiota in elderly pneumonia patients. ISME J 10, 97–108 (2016).

14. Chanderraj, R. et al. In critically ill patients, anti-anaerobic antibiotics increase risk of adverse clinical outcomes. Eur Respir J 61, 2200910 (2023).

15. Dickson, R. P. et al. Spatial Variation in the Healthy Human Lung Microbiome and the Adapted Island Model of Lung Biogeography. Ann Am Thorac Soc 12, 821–830 (2015).

16. Edouard, S. et al. The nasopharyngeal microbiota in patients with viral respiratory tract infections is enriched in bacterial pathogens. Eur J Clin Microbiol Infect Dis 37, 1725–1733 (2018).

17. Zelasko, S. et al. Early-life upper airway microbiota are associated with decreased lower respiratory tract infections. J Allergy Clin Immunol 155, 436–450 (2025).

18. Ramsheh, M. Y. et al. Lung microbiome composition and bronchial epithelial gene expression in patients with COPD versus healthy individuals: a bacterial 16S rRNA gene sequencing and host transcriptomic analysis. Lancet Microbe 2, e300–e310 (2021).

19. Edouard, S. et al. The nasopharyngeal microbiota in patients with viral respiratory tract infections is enriched in bacterial pathogens. Eur J Clin Microbiol Infect Dis 37, 1725–1733 (2018).

20. Bousbia, S. et al. Repertoire of intensive care unit pneumonia microbiota. PLoS One 7, e32486 (2012).

21. Melo-Dias, S. et al. Responsiveness to pulmonary rehabilitation in COPD is associated with changes in microbiota. Respir Res 24, 29 (2023).

22. Segal, L. N. et al. Enrichment of lung microbiome with supraglottic taxa is associated with increased pulmonary inflammation. Microbiome 1, 19 (2013).

23. Segal, L. N. et al. Enrichment of the lung microbiome with oral taxa is associated with lung inflammation of a Th17 phenotype. Nat Microbiol 1, 16031 (2016).

24. Das, S. et al. A prevalent and culturable microbiota links ecological balance to clinical stability of the human lung after transplantation. Nat Commun 12, 2126 (2021).

25. Wu, B. G. et al. Episodic Aspiration with Oral Commensals Induces a MyD88-dependent, Pulmonary T-Helper Cell Type 17 Response that Mitigates Susceptibility to Streptococcus pneumoniae. Am J Respir Crit Care Med 203, 1099–1111 (2021).

26. Horn, K. J., Schopper, M. A., Drigot, Z. G. & Clark, S. E. Airway Prevotella promote TLR2-dependent neutrophil activation and rapid clearance of Streptococcus pneumoniae from the lung. Nat Commun 13, 3321 (2022).

27. Goryachok, M. et al. Functional CFTR may be required for Prevotella melaninogenica regulation of epithelial cell defense against Staphylococcus aureus. J Cyst Fibros 25, 340–352 (2026).

28. Barletta, K. E. et al. Leukocyte compartments in the mouse lung: distinguishing between marginated, interstitial, and alveolar cells in response to injury. J Immunol Methods 375, 100–110 (2012).

29. Mould, K. J. et al. Cell Origin Dictates Programming of Resident versus Recruited Macrophages during Acute Lung Injury. Am J Respir Cell Mol Biol 57, 294–306 (2017).

30. Mould, K. J., Jackson, N. D., Henson, P. M., Seibold, M. & Janssen, W. J. Single cell RNA sequencing identifies unique inflammatory airspace macrophage subsets. JCI Insight 4, e126556, 126556 (2019).

31. Hou, F. et al. Distinct Transcriptional and Functional Differences of Lung Resident and Monocyte-Derived Alveolar Macrophages During the Recovery Period of Acute Lung Injury. Immune Netw 23, e24 (2023).

32. Gordon, D. L., Rice, J. L. & McDonald, P. J. Regulation of human neutrophil type 3 complement receptor (iC3b receptor) expression during phagocytosis of Staphylococcus aureus and Escherichia coli. Immunology 67, 460–465 (1989).

33. Prince, L. R., Whyte, M. K., Sabroe, I. & Parker, L. C. The role of TLRs in neutrophil activation. Curr Opin Pharmacol 11, 397–403 (2011).

34. Lawrence, S. M., Corriden, R. & Nizet, V. The Ontogeny of a Neutrophil: Mechanisms of Granulopoiesis and Homeostasis. Microbiol Mol Biol Rev 82, e00057–17 (2018).

35. Borregaard, N., Sørensen, O. E. & Theilgaard-Mönch, K. Neutrophil granules: a library of innate immunity proteins. Trends Immunol 28, 340–345 (2007).

36. Faustman, D. & Davis, M. TNF receptor 2 pathway: drug target for autoimmune diseases. Nat Rev Drug Discov 9, 482–493 (2010).

37. Parker, D. et al. Streptococcus pneumoniae DNA initiates type I interferon signaling in the respiratory tract. mBio 2, e00016–00011 (2011).

38. Maier, B. B. et al. Type I interferon promotes alveolar epithelial type II cell survival during pulmonary Streptococcus pneumoniae infection and sterile lung injury in mice. Eur J Immunol 46, 2175–2186 (2016).

39. Damjanovic, D. et al. Type 1 interferon gene transfer enhances host defense against pulmonary Streptococcus pneumoniae infection via activating innate leukocytes. Mol Ther Methods Clin Dev 1, 5 (2014).

40. Maier, B. B. et al. Type I interferon promotes alveolar epithelial type II cell survival during pulmonary Streptococcus pneumoniae infection and sterile lung injury in mice. Eur J Immunol 46, 2175–2186 (2016).

41. Li, W., Moltedo, B. & Moran, T. M. Type I interferon induction during influenza virus infection increases susceptibility to secondary Streptococcus pneumoniae infection by negative regulation of γδ T cells. J Virol 86, 12304–12312 (2012).

42. Shahangian, A. et al. Type I IFNs mediate development of postinfluenza bacterial pneumonia in mice. J Clin Invest 119, 1910–1920 (2009).

43. Nakamura, S., Davis, K. M. & Weiser, J. N. Synergistic stimulation of type I interferons during influenza virus coinfection promotes Streptococcus pneumoniae colonization in mice. J Clin Invest 121, 3657–3665 (2011).

44. Ivashkiv, L. B. & Donlin, L. T. Regulation of type I interferon responses. Nat Rev Immunol 14, 36–49 (2014).

45. Carow, B. & Rottenberg, M. E. SOCS3, a Major Regulator of Infection and Inflammation. Front Immunol 5, 58 (2014).

46. Kawai, T. & Akira, S. The role of pattern-recognition receptors in innate immunity: update on Toll-like receptors. Nat Immunol 11, 373–384 (2010).

47. Parsa, K. V. L., Butchar, J. P., Rajaram, M. V. S., Cremer, T. J. & Tridandapani, S. The tyrosine kinase Syk promotes phagocytosis of Francisella through the activation of Erk. Mol Immunol 45, 3012–3021 (2008).

48. Lavoie, H., Gagnon, J. & Therrien, M. ERK signalling: a master regulator of cell behaviour, life and fate. Nat Rev Mol Cell Biol 21, 607–632 (2020).

49. Shi, Y. et al. Protein-tyrosine kinase Syk is required for pathogen engulfment in complement-mediated phagocytosis. Blood 107, 4554–4562 (2006).

50. Koppe, U. et al. Streptococcus pneumoniae stimulates a STING- and IFN regulatory factor 3-dependent type I IFN production in macrophages, which regulates RANTES production in macrophages, cocultured alveolar epithelial cells, and mouse lungs. J Immunol 188, 811–817 (2012).

51. Lee, B. et al. Influenza-induced type I interferon enhances susceptibility to gram-negative and gram-positive bacterial pneumonia in mice. Am J Physiol Lung Cell Mol Physiol 309, L158–167 (2015).

52. Standish, A. J. & Weiser, J. N. Human neutrophils kill Streptococcus pneumoniae via serine proteases. J Immunol 183, 2602–2609 (2009).

53. Herold, S. et al. Lung epithelial apoptosis in influenza virus pneumonia: the role of macrophage-expressed TNF-related apoptosis-inducing ligand. J Exp Med 205, 3065–3077 (2008).

54. Traber, K. E. & Mizgerd, J. P. The Integrated Pulmonary Immune Response to Pneumonia. Annu Rev Immunol 43, 545–569 (2025).

55. Pernet, E., Downey, J., Vinh, D. C., Powell, W. S. & Divangahi, M. Leukotriene B4-type I interferon axis regulates macrophage-mediated disease tolerance to influenza infection. Nat Microbiol 4, 1389–1400 (2019).

56. Xiao, K. et al. A pan-immune panorama of bacterial pneumonia revealed by a large-scale single-cell transcriptome atlas. Signal Transduct Target Ther 10, 5 (2025).

57. Xiao, K. et al. A Large-Scale Single-Cell Atlas Reveals the Peripheral Immune Panorama of Bacterial Pneumonia. Am J Respir Crit Care Med 211, 2363–2381 (2025).

58. Guillon, A. et al. Pneumonia recovery reprograms the alveolar macrophage pool. JCI Insight 5, e133042, 133042 (2020).

59. Ngo, V. L., et al. Segmented filamentous bacteria reprogramming of alveolar macrophages limits postinfluenza bacterial pneumonia. Sci Immunol 11, eadt8858 (2026).

60. Chakraborty, S. et al. Trained immunity of alveolar macrophages enhances injury resolution via KLF4-MERTK-mediated efferocytosis. J Exp Med 220, e20221388 (2023).

61. Zahalka, S. et al. Trained immunity of alveolar macrophages requires metabolic rewiring and type 1 interferon signaling. Mucosal Immunol 15, 896–907 (2022).

62. Theobald, H. et al. Apolipoprotein E controls Dectin-1-dependent development of monocyte-derived alveolar macrophages upon pulmonary β-glucan-induced inflammatory adaptation. Nat Immunol 25, 994–1006 (2024).

63. Xie, X. et al. Single-cell transcriptome profiling reveals neutrophil heterogeneity in homeostasis and infection. Nat Immunol 21, 1119–1133 (2020).

64. Cortjens, B. et al. Neutrophil subset responses in infants with severe viral respiratory infection. Clin Immunol 176, 100–106 (2017).

65. Matarazzo, L. et al. Neutrophil subsets enhance the efficacy of host-directed therapy in pneumococcal pneumonia. Mucosal Immunol 18, 257–268 (2025).

66. Grell, M. et al. The transmembrane form of tumor necrosis factor is the prime activating ligand of the 80 kDa tumor necrosis factor receptor. Cell 83, 793–802 (1995).

67. Eissner, G., Kolch, W. & Scheurich, P. Ligands working as receptors: reverse signaling by members of the TNF superfamily enhance the plasticity of the immune system. Cytokine Growth Factor Rev 15, 353–366 (2004).

68. Miralda, I., Uriarte, S. M. & McLeish, K. R. Multiple Phenotypic Changes Define Neutrophil Priming. Front Cell Infect Microbiol 7, 217 (2017).

69. Youn, C. et al. Neutrophil-intrinsic TNF receptor signaling orchestrates host defense against Staphylococcus aureus. Sci Adv 9, eadf8748 (2023).

70. Hahn, I. et al. Cathepsin G and neutrophil elastase play critical and nonredundant roles in lung-protective immunity against Streptococcus pneumoniae in mice. Infect Immun 79, 4893–4901 (2011).

71. Standish, A. J. & Weiser, J. N. Human neutrophils kill Streptococcus pneumoniae via serine proteases. J Immunol 183, 2602–2609 (2009).

72. Netea, M. G. et al. Defining trained immunity and its role in health and disease. Nat Rev Immunol 20, 375–388 (2020).

73. Moorlag, S. J. C. F. M. et al. BCG Vaccination Induces Long-Term Functional Reprogramming of Human Neutrophils. Cell Rep 33, 108387 (2020).

74. Mitroulis, I. et al. Modulation of Myelopoiesis Progenitors Is an Integral Component of Trained Immunity. Cell 172, 147–161.e12 (2018).

75. Kalafati, L. et al. Innate Immune Training of Granulopoiesis Promotes Anti-tumor Activity. Cell 183, 771–785.e12 (2020).

76. Ajit, J. et al. β-glucan induced trained immunity enhances antibody levels in a vaccination model in mice. PLoS One 20, e0323376 (2025).

77. Resiliac, J., Rohlfing, M., Santoro, J., Hussain, S.-R. A. & Grayson, M. H. Low-Dose Lipopolysaccharide Protects from Lethal Paramyxovirus Infection in a Macrophage-and TLR4-Dependent Process. J Immunol 210, 348–355 (2023).

78. Wong, K. K. et al. Microbial contribution to metabolic niche formation varies across the respiratory tract. Cell Host Microbe 33, 1073–1088.e6 (2025).

79. Chanderraj, R. et al. Mortality of Patients With Sepsis Administered Piperacillin-Tazobactam vs Cefepime. JAMA Intern Med 184, 769–777 (2024).

80. Kullberg, R. F. J., Schinkel, M. & Wiersinga, W. J. Empiric anti-anaerobic antibiotics are associated with adverse clinical outcomes in emergency department patients. Eur Respir J 61, 2300413 (2023).

81. Kullberg, R. F. J. et al. Empirical antibiotic therapy for sepsis: save the anaerobic microbiota. Lancet Respir Med 13, 92–100 (2025).

82. Kitsios, G. D. et al. Longitudinal multicompartment characterization of host-microbiota interactions in patients with acute respiratory failure. Nat Commun 15, 4708 (2024).

83. Zafar, M. A., Kono, M., Wang, Y., Zangari, T. & Weiser, J. N. Infant Mouse Model for the Study of Shedding and Transmission during Streptococcus pneumoniae Monoinfection. Infect Immun 84, 2714–2722 (2016).

84. Mack, M. et al. Expression and characterization of the chemokine receptors CCR2 and CCR5 in mice. J Immunol 166, 4697–4704 (2001).

85. Wolock, S. L., Lopez, R. & Klein, A. M. Scrublet: Computational Identification of Cell Doublets in Single-Cell Transcriptomic Data. Cell Syst 8, 281–291.e9 (2019).

86. Quan, F. et al. Annotation of cell types (ACT): a convenient web server for cell type annotation. Genome Med 15, 91 (2023).

87. Wolf, F. A. et al. PAGA: graph abstraction reconciles clustering with trajectory inference through a topology preserving map of single cells. Genome Biol 20, 59 (2019).

88. Wu, T. et al. clusterProfiler 4.0: A universal enrichment tool for interpreting omics data. Innovation (Camb) 2, 100141 (2021).

89. Jin, S., Plikus, M. V. & Nie, Q. CellChat for systematic analysis of cell-cell communication from single-cell transcriptomics. Nat Protoc 20, 180–219 (2025).

90. Jacomy, M., Venturini, T., Heymann, S. & Bastian, M. ForceAtlas2, a continuous graph layout algorithm for handy network visualization designed for the Gephi software. PLoS One 9, e98679 (2014).

