## Supplemental Information for "Myeloid cell reprogramming drives enhanced defense against *Streptococcus pneumoniae* lung infection following exposure to commensal *Prevotella*"

### Supplementary Data

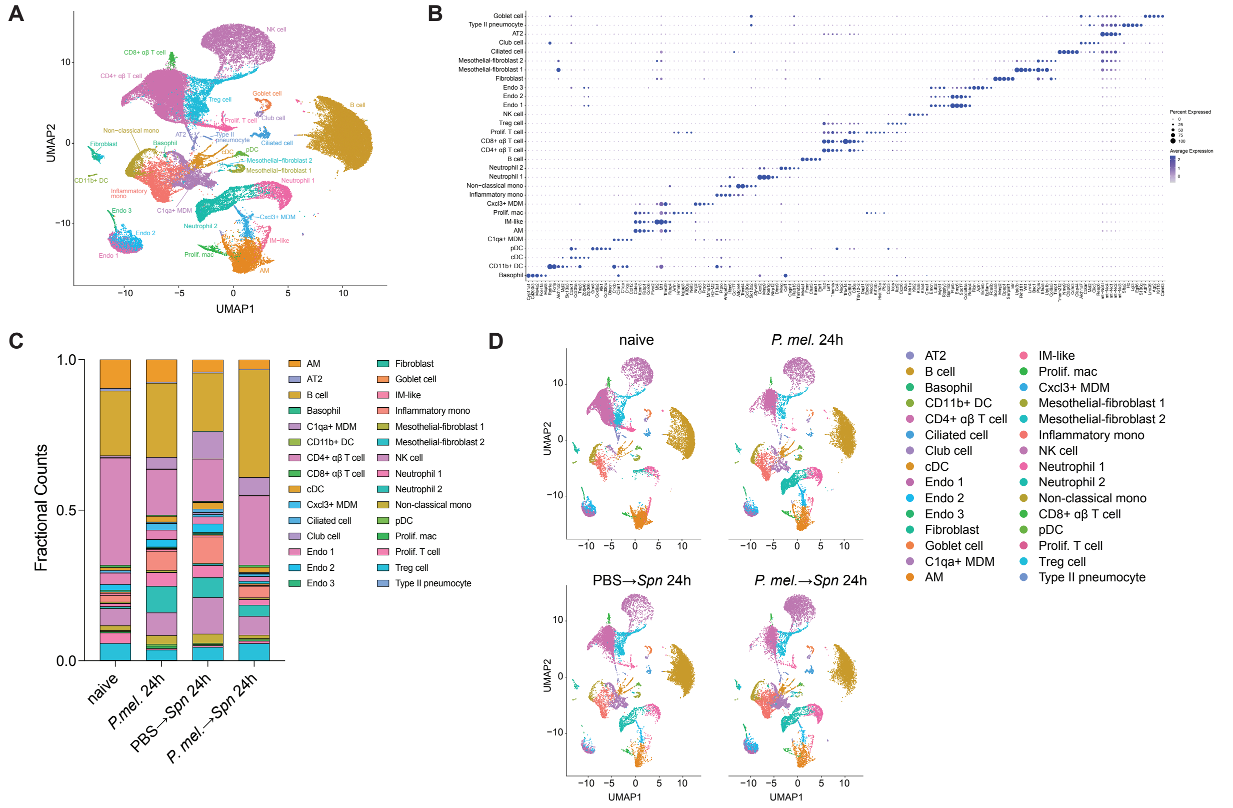


**Figure S1: Landscape of lung tissue cellular response to *P. melaninogenica* and *Spn* infection detected by scRNA-seq.**

(**A**) UMAP projection displaying all lung tissue cell populations identified by scRNA-seq. (**B**) Top 5 DEGs for all cell populations. (**C**) Fractional counts for all cell populations split by condition. (**D**) UMAP projections of all cell populations split by condition. Abbreviations: AT2 = alveolar type-2, endo = endothelial, NK = natural killer, Treg = T regulatory cell, Prolif = proliferating, mono = monocyte, MDM = monocyte-derived macrophage, IM = interstitial macrophage, mac = macrophage, AM = alveolar macrophage, DC = dendritic cell, pDC = plasmacytoid DC, cDC = conventional DC.

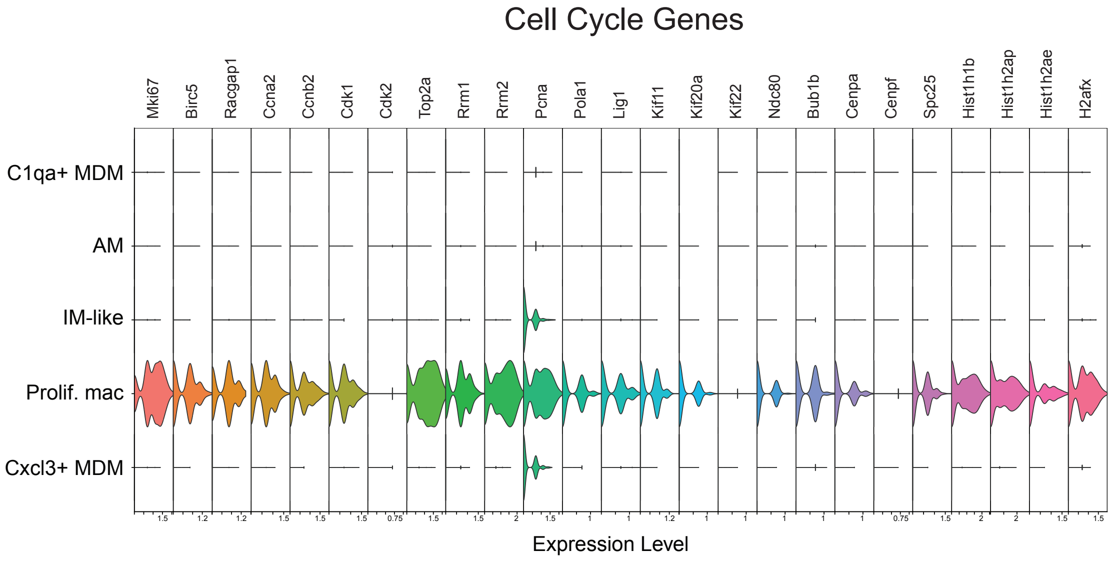


**Figure S2: Expression of cell cycle genes in macrophages.**

Violin plot showing expression of 25 cell cycle genes across macrophage populations. Abbreviations: Prolif = proliferating, mono = monocyte, MDM = monocyte-derived macrophage, IM = interstitial macrophage, mac = macrophage, AM = alveolar macrophage.

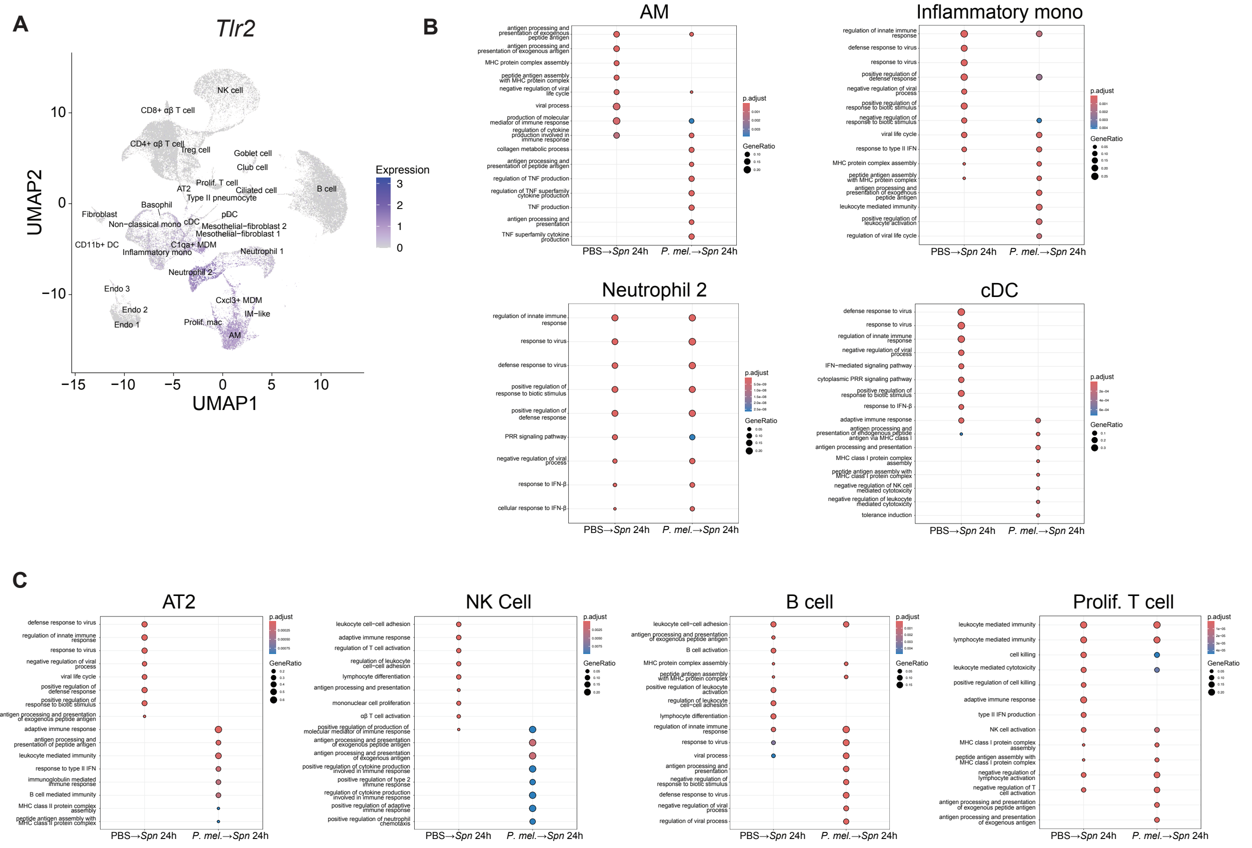


**Figure S3: *P. melaninogenica* priming upregulates distinct immune pathways.** (**A**) UMAP overlay of *Tlr2* gene expression. Unbiased pathway analysis indicating the top gene ontology (GO) terms for (**B**) selected myeloid cell populations and (**C**) select non-myeloid populations in *Spn*-infected mice with or without i.t. exposure to *P. mel* HK 24 hours prior to infection. Abbreviations: AM = alveolar macrophage, mono = monocyte, cDC = conventional DC, AT2 = alveolar type-2, NK = natural killer, Prolif. = proliferating.

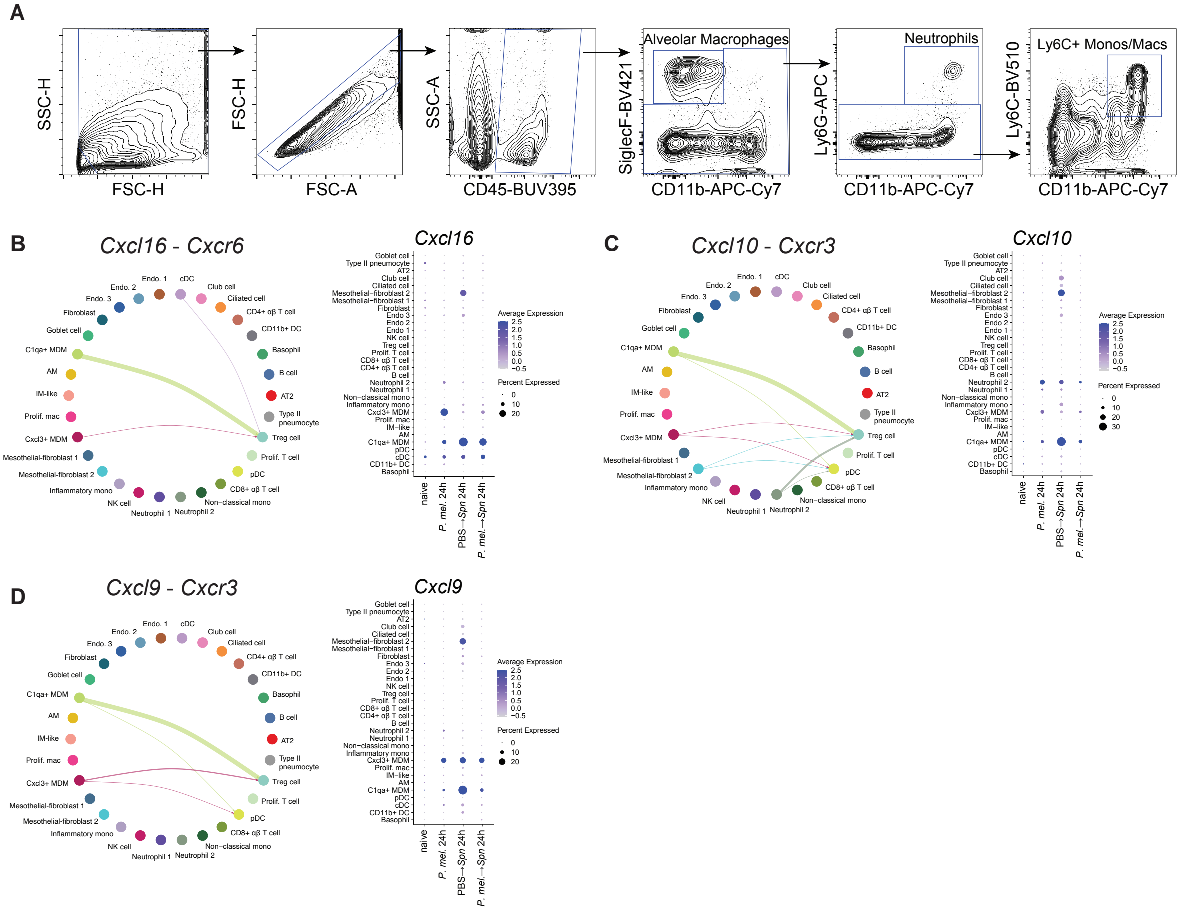
A
SSC-H
FSC-H
FSC-H
FSC-A
SSC-A
CD45-BUV395
Alveolar Macrophages
SiglecF-BV421
CD11b-APC-Cy7
Neutrophils
Ly6G-APC
CD11b-APC-Cy7
Ly6C+ Monos/Macs
Ly6C-BV510
CD11b-APC-Cy7
B
Cxcl16 - Cxcr6
Cxcl16
C
Cxcl10 - Cxcr3
Cxcl10
D
Cxcl9 - Cxcr3
Cxcl9
Endo. 1
cDC
Club cell
Ciliated cell
CD4+ αβ T cell
CD11b+ DC
Basophil
B cell
AT2
Type II pneumocyte
Treg cell
Prolif. T cell
pDC
CD8+ αβ T cell
Non-classical mono
Neutrophil 2
Neutrophil 1
NK cell
Inflammatory mono
Mesothelial-fibroblast 2
Mesothelial-fibroblast 1
Cxcl3+ MDM
Prolif. mac
IM-like
AM
C1qa+ MDM
Goblet cell
Fibroblast
Endo. 3
Endo. 2
Average Expression
Percent Expressed
naive
P. mel. 24h
PBS→Spn 24h
P. mel.→Spn 24h

**Figure S4: Macrophages signal to Tregs through chemokine expression.**

(**A**) Flow cytometry gating strategy to identify AMs, neutrophils, and Ly6C+ monos/macs. Gating strategy was used for all flow cytometry experiments. CellChat circle plots of inferred cell-cell communication networks for *Cxcl16*-*Cxcr6* (**B**) *Cxcl10*-*Cxcr3* (**C**), and *Cxcr9*-*Cxcr3* (**D**) signaling along with dot plots showing expression of corresponding chemokine genes by condition. Abbreviations: AT2 = alveolar type-2, endo = endothelial, NK = natural killer, Treg = T regulatory cell, Prolif = proliferating, mono = monocyte, MDM = monocyte-derived macrophage, IM = interstitial macrophage, mac = macrophage, AM = alveolar macrophage, DC = dendritic cell, pDC = plasmacytoid DC, cDC = conventional DC.

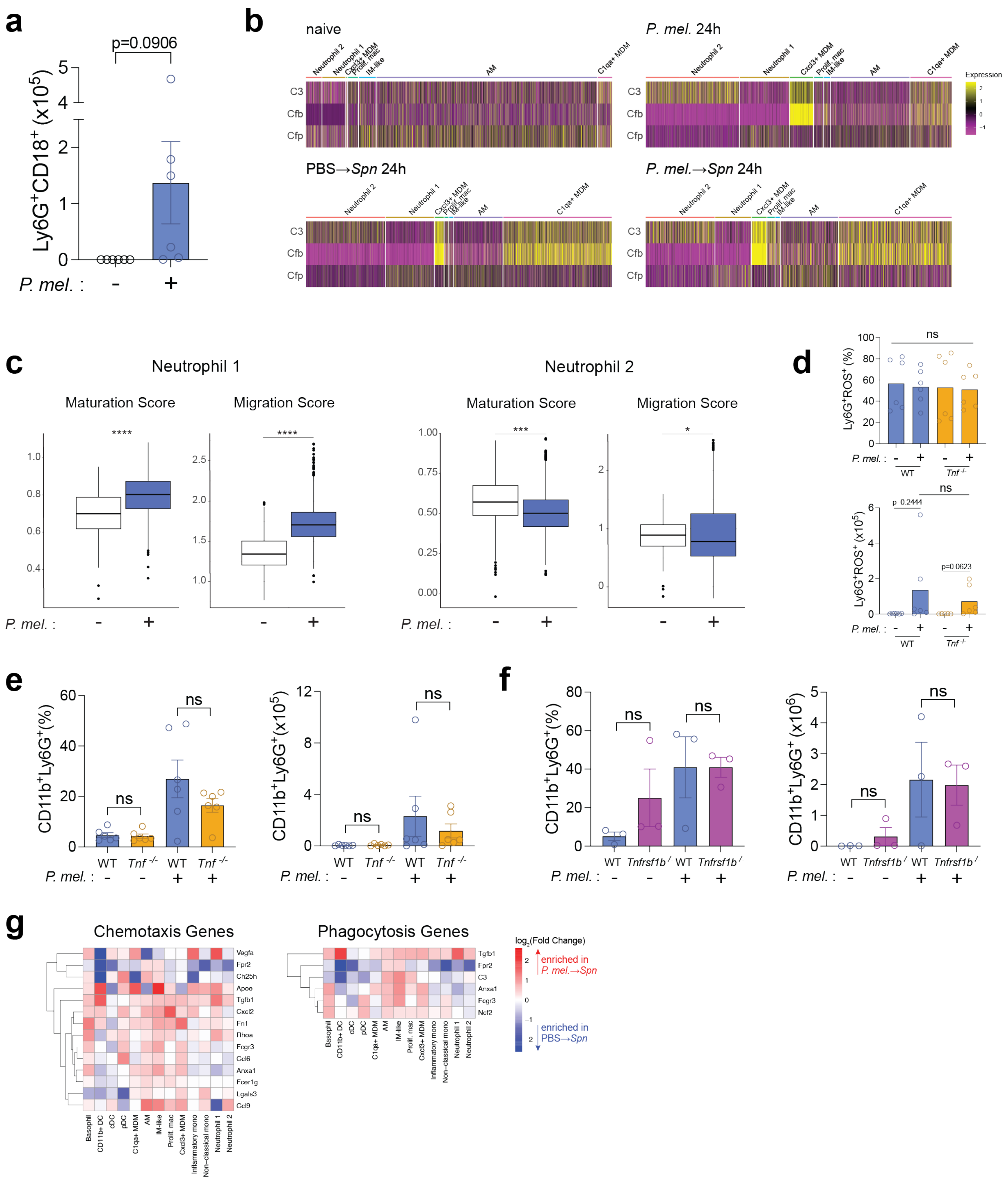

a
p=0.0906
Ly6G+CD18+ (x10^5)
P. mel. :
b
naive
P. mel. 24h
PBS→Spn 24h
P. mel.→Spn 24h
C3
Cfb
Cfp
Neutrophil 2
Neutrophil 1
Cxcl3+ MDM
Prolif. mac
IM-like
AM
C1qa+ MDM
Expression
c
Neutrophil 1
Neutrophil 2
Maturation Score
Migration Score
P. mel. :
d
Ly6G+ROS+ (%)
Ly6G+ROS+ (x10^5)
ns
p=0.2444
p=0.0623
WT
Tnf -/-
e
CD11b+Ly6G+(%)
CD11b+Ly6G+(x10^5)
f
CD11b+Ly6G+(%)
CD11b+Ly6G+ (x10^6)
Tnfrsf1b-/-
g
Chemotaxis Genes
Phagocytosis Genes
Vegfa
Fpr2
Ch25h
Apoe
Tgfb1
Cxcl2
Fn1
Rhoa
Fcgr3
Ccl6
Anxa1
Fcer1g
Lgals3
Ccl9
C3
Ncf2
Basophil
CD11b+ DC
cDC
pDC
C1qa+ MDM
AM
IM-like
Prolif. mac
Cxcl3+ MDM
Inflammatory mono
Non-classical mono
Neutrophil 1
Neutrophil 2
log2(Fold Change)
enriched in P. mel.→Spn
enriched in PBS→Spn

**Figure S5: TNF signaling does not regulate neutrophil ROS production during *P. melaninogenica* exposure.**

(**A**) Quantification of Ly6G+CD18+ neutrophil cell numbers in lung tissue of mice exposed to FITC-labeled *Spn* HK for 3 hours, with prior exposure to PBS or *P. mel* HK (mean ± SEM; n = 6 biological replicates pooled from 2 independent experiments). (**B**) Heatmap displaying alternative complement pathway gene expression across macrophage and neutrophil populations, split by condition. (**C**) Module scoring analysis of maturation and migration signature genes in neutrophil populations of naïve mice and mice exposed to *P. mel* HK. (**D**) Quantification of Ly6G+ROS+ neutrophils via flow cytometry in WT and $Tnf^{-/-}$ mice along with 24-hour exposure to PBS or *P. mel* HK (mean ± SEM; n = 6 biological replicates pooled from 2 independent experiments). Quantification of CD11b+Ly6G+ neutrophils via flow cytometry in WT vs $Tnf^{-/-}$ (**E**) or $Tnfrsf1b^{-/-}$ (**F**) mice exposed to PBS or *P. mel* HK (mean ± SEM; n = 3-6 biological replicates pooled from 1-2 experiments). (**G**) Heatmaps displaying log-fold changes of chemotaxis and phagocytosis gene expression in myeloid cells of mice exposed to *P. mel* HK prior to *Spn* infection, relative to mice infected with *Spn* alone. Abbreviations: AT2 = alveolar type-2, endo = endothelial, NK = natural killer, Treg = T regulatory cell, Prolif = proliferating, mono = monocyte, MDM = monocyte-derived macrophage, IM = interstitial macrophage, mac = macrophage, AM = alveolar macrophage, DC = dendritic cell, pDC = plasmacytoid DC, cDC = conventional DC. Data were analyzed by unpaired t test (**A**), Wilcoxon rank-sum test (**C**), or Kruskal-Wallis with Dunn's post hoc test (**D,E,F**). *$p$<0.05, ***$p$<0.001, ****$p$<0.0001.

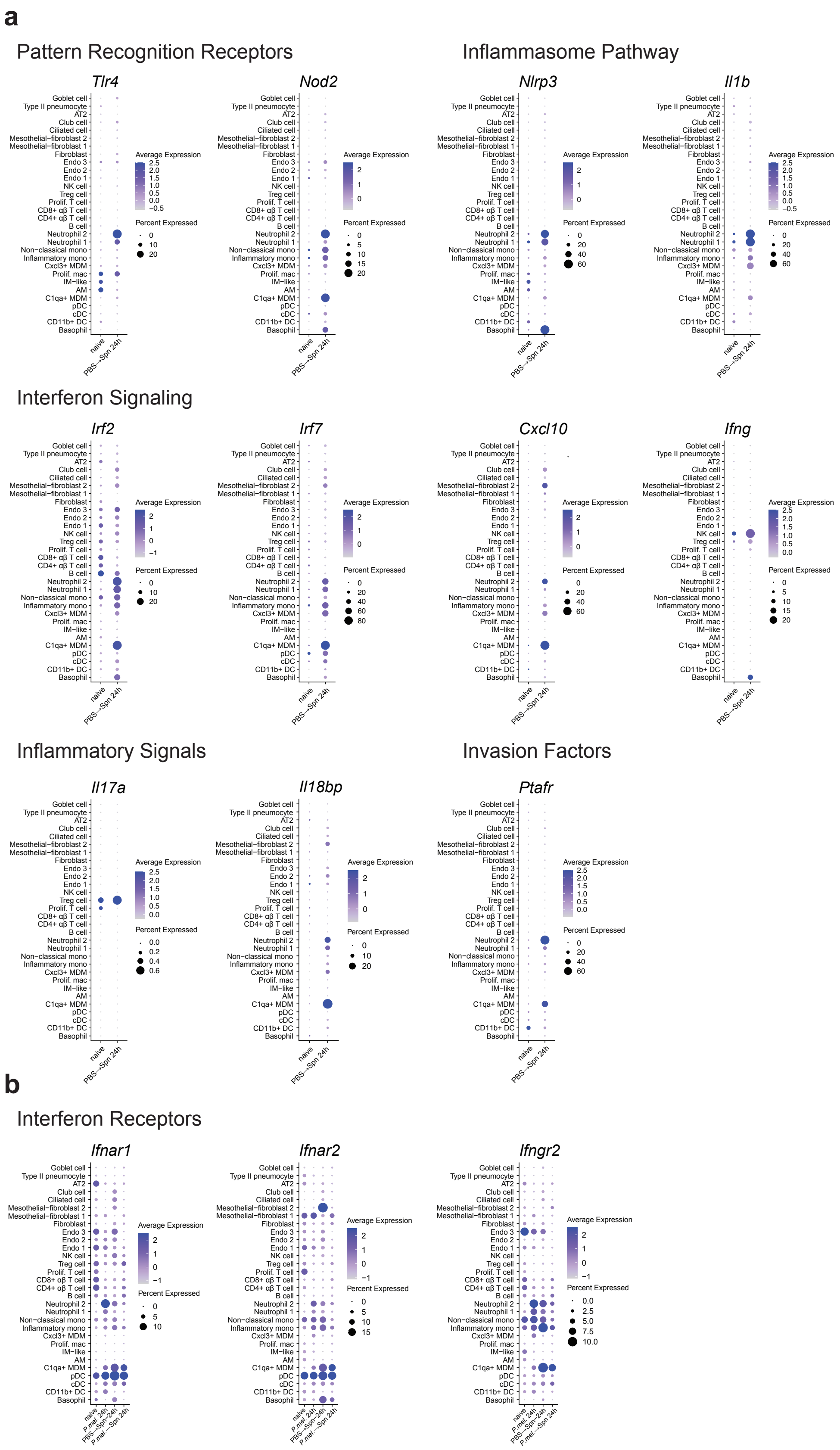
a
Pattern Recognition Receptors
Tlr4
Nod2
Inflammasome Pathway
Nlrp3
Il1b
Interferon Signaling
Irf2
Irf7
Cxcl10
Ifng
Inflammatory Signals
Il17a
Il18bp
Invasion Factors
Ptafr
b
Interferon Receptors
Ifnar1
Ifnar2
Ifngr2
Average Expression
Percent Expressed
naive
PBS→Spn 24h
P.mel. 24h
P.mel.→Spn 24h
Goblet cell
Type II pneumocyte
AT2
Club cell
Ciliated cell
Mesothelial−fibroblast 2
Mesothelial−fibroblast 1
Fibroblast
Endo 3
Endo 2
Endo 1
NK cell
Treg cell
Prolif. T cell
CD8+ αβ T cell
CD4+ αβ T cell
B cell
Neutrophil 2
Neutrophil 1
Non−classical mono
Inflammatory mono
Cxcl3+ MDM
Prolif. mac
IM−like
AM
C1qa+ MDM
pDC
cDC
CD11b+ DC
Basophil

**Figure S6: *S. pneumoniae* infection induces expression of canonical genes involved in host response to infection.**

(**A**) Dot plots showing expression of genes involved in pattern recognition receptor signaling, inflammasome signaling, interferon signaling, inflammatory signaling, and host cell adhesion across all cell populations in naïve vs *Spn*-infected mice. (**B**) Dot plots showing expression of interferon receptor genes across all cell populations and all conditions. Abbreviations: AT2 = alveolar type-2, endo = endothelial, NK = natural killer, Treg = T regulatory cell, Prolif = proliferating, mono = monocyte, MDM = monocyte-derived macrophage, IM = interstitial macrophage, mac = macrophage, AM = alveolar macrophage, DC = dendritic cell, pDC = plasmacytoid DC, cDC = conventional DC.

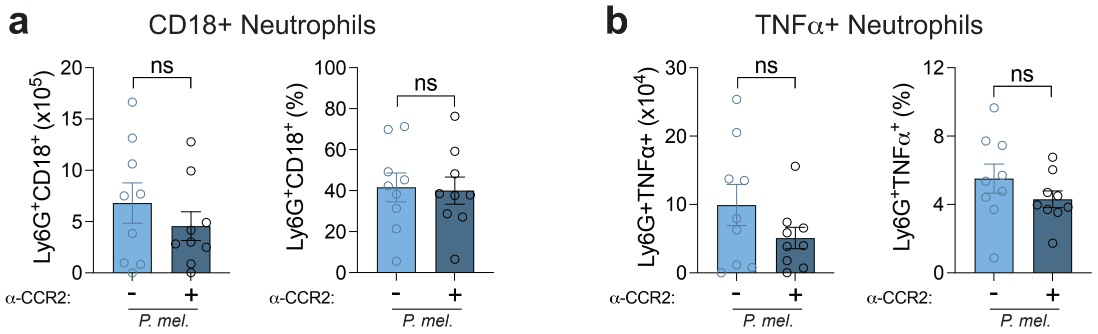


**Figure S7: CCR2+ depletion does not impact neutrophil activation after *P. melaninogenica* exposure.**

Flow cytometry quantification of (**A**) Ly6G+CD18+ neutrophils and (**B**) TNFα+ neutrophils in mice treated with isotype control or anti-CCR2 antibodies prior to *P. mel* HK exposure (mean ± SEM; n = 9 biological replicates pooled from 3 independent experiments). Data were analyzed by unpaired t test.
